# Robust analytical methods for bis(monoacylglycero)phosphate profiling in health and disease

**DOI:** 10.1101/2025.02.13.638174

**Authors:** Wentao Dong, Kwamina Nyame, Eshaan S. Rawat, Uche N. Medoh, Jian Xiong, Caio C. Bonin, Hisham N. Alsohybe, Hunter Y. Liu, Sara Gomes, Tammy Hsieh, Marianna Arnold, Frank Hsieh, Esther Sammler, Monther Abu-Remaileh

**Affiliations:** Department of Chemical Engineering, Stanford University, Stanford, CA, 94305, USA; Department of Genetics, Stanford University, Stanford, CA, 94305, USA; The Institute for Chemistry, Engineering & Medicine for Human Health (Sarafan ChEM-H), Stanford University, Stanford, CA, 94305, USA; Aligning Science Across Parkinson’s (ASAP) Collaborative Research Network, Chevy Chase, MD, 20815, USA; Department of Biochemistry, Stanford University, Stanford, CA 94305, USA; Arc Institute, Palo Alto, CA 94304, USA; Department of Bioengineering, Stanford University, Stanford, CA, 94305, USA; Stanford Biophysics Program, Stanford University, Stanford, CA 94305, USA; Medical Research Council Protein Phosphorylation and Ubiquitylation Unit, University of Dundee, Dundee DD1 5EH, UK; Nextcea, Inc., Woburn, MA, USA; The Phil & Penny Knight Initiative for Brain Resilience at the Wu Tsai Neurosciences Institute, Stanford University, Stanford, CA 94305, USA

**Keywords:** Liquid chromatography/mass spectrometry (LC/MS), Lysosome, Lipid metabolism, Bis(monoacylglycero)phosphate (BMP), Phosphatidylglycerol (PG), Lysobisphosphatidic acid (LBPA), Triple quadrupole, Orbitrap, CLN5, Neurodegeneration

## Abstract

Bis(monoacylglycero)phosphate (BMP), a distinct anionic phospholipid predominantly found in late endosomes and lysosomes, plays a pivotal role in supporting lysosomal functions and maintaining metabolic homeostasis. Its impaired function is associated with an array of disorders, notably neurodegenerative diseases. However, the identification and quantitation of BMP remains difficult due to its structural similarity to isomer phosphatidylglycerol (PG), thus necessitating robust analytical methods for accurate and reliable BMP profiling. In this study, we present comprehensive liquid chromatography – tandem mass spectrometry (MS2) methodologies for the precise and systematic analysis of BMP species in biological samples. We detail LC/MS methods for both an untargeted Orbitrap mass spectrometer and a targeted triple quadrupole (QQQ) mass spectrometer. We utilize differences in polarity and structure to annotate BMPs and PGs based on retention time and positive mode MS2 fragmentation patterns, respectively. Further, we propose a new approach for overcoming common challenges in BMP profiling by leveraging the newly discovered biochemical function of CLN5 as the BMP synthase. Since genetic ablation of CLN5 leads to specific depletion of BMPs but not PGs, we use lipid extracts from *CLN5* knockout (KO) and wild-type (WT) cells as biological standards to confidently annotate BMPs as targets with significantly low BMP Identification Index (BMPII), defined as BMPII = *CLN5* KO / WT. We additionally propose the BMP enrichment score (BMPES) as a secondary validation metric, defined as lysosomal abundance of BMP / whole-cell abundance. Altogether, this approach constitutes a robust method for BMP profiling and annotation, furthering research into health and disease.

## Introduction

Bis(monoacylglycero)phosphate (BMP), also mistakenly known as lysobisphosphatidic acid (LBPA), is a unique lipid class characterized by two monoglycerides linked via a central phosphate group^1,2^. BMPs make up less than 1% of the total lipid composition of most cells and tissues^1,3–5^. However, they are specifically localized to the inner membranes of the late endosome and lysosome, where they play a critical role in lysosomal function^6,7^. In the late endosome and lysosome, BMPs comprise up to 16% of the total lipid content^1,4,8^, as well as up to 77% of the lipid content of intraluminal vesicles (ILVs) within the lysosome^8^. Of the BMPs found within the endocytic pathway, only around 5% are found in the late endosome, with the remaining 95% found within the lysosome^9^. BMP holds a negative charge within the acidic lysosome, while proteins frequently gain a positive charge within the lysosome. This allows BMP to act as a docking site and cofactor for a variety of intraluminal lysosomal proteins^1,10^. For instance, the breakdown of glucosylceramides by acid ß-glucosidase encoded by *GBA* (GCase) with the help of its saposin C activator protein (Sap-C) relies upon both GCase and Sap-C binding to BMPs on ILVs^11^. Hydrolysis of sphingomyelin by acid sphingomyelinase (ASM) likewise relies on BMP-mediated anchoring of ASM to the ILVs^12^. Lysosomal phospholipase A2 and saposins A through D are some of the other proteins that have been shown to interact with BMP^13,14^. In addition, BMP indirectly affects lysosomal stability and lipid homeostasis by binding to heat shock protein 70 (Hsp70)^15^. At the acidic pH of the lysosome, the ATPase domain of Hsp70 becomes positively charged and binds to BMP, thus stabilizing lysosomal membranes.

Given their prominent role in lysosomal homeostasis, BMPs are unsurprisingly dysregulated in a variety of lysosomal storage disorders and other diseases^1,10,16^. Gaucher’s disease, one of the most common lysosomal storage disorders resulting from a loss of function of the gene encoding acid ß-glucosidase, has been linked to BMP accumulation^16,17^. Additionally, BMPs accumulate in the urine of patients with Niemann-Pick disease^18^. Outside of lysosomal storage disorders, BMP accumulates in the brains of patients with mixed dementia^19^ and Alzheimer’s disease^20^. Similarly, a deficiency in mice of Cathepsin D, a protease that degrades the amyloid ß-protein and tau protein and has been implicated in Alzheimer’s disease, leads to BMP accumulation in the brain^21,22^. Further, elevated BMP has also been found in the urine of patients with varying mutations associated with Parkinson’s disease^23–25^. This clinically significant presence of BMP has led to urine BMP levels being used as an interventional biomarker in LRRK2 inhibitor clinical trials for Parkinson’s disease^26,27^. Another link to BMP has been found by studying the frontotemporal dementia gene *GRN* (*CLN11*). Deficiency of progranulin (PGRN) has been shown to markedly reduce BMP levels, leading to impaired ganglioside and glucosylceramide catabolism^28,29^. Similarly, reductions in BMP levels have also been observed in CLN3 Batten disease^30–32^. Finally, BMP levels have been shown to be altered in mouse and human aging and can be reversed by exercise^33,34^.

While BMP levels are altered in several human conditions and targeting the BMP pathway holds immense therapeutic promise^6^, accurate quantitation of BMP species remains limited to few highly specialized laboratories, thus impeding progress in this quickly emerging field. BMP is a structural isomer of phosphatidylglycerol (PG), making separation of the two lipid classes difficult across a wide variety of detection methods. Early methods for BMP profiling involved the use of monoclonal antibodies against BMP, which, when combined with immunofluorescence, allow for quantitation of total cellular BMP content^35^. This method has been used successfully to study BMP, and was recently used in combination with flow cytometry to screen for regulators of cholesterol and BMP^36^. However, these antibodies cannot distinguish between individual BMPs with different side chain compositions, and lack the quantitative power of analytical methods such as mass spectrometry. Another method to profile BMP involves using a pseudoisocyanine dye that forms J-aggregates when bound to BMP and can be visualized directly with confocal microscopy^37^; however, the dye has the same lack of specificity and quantitative power as antibody-based techniques. Though BMP and PG are structural isomers, past research has made progress towards distinguishing the two with LC/MS-based techniques. Methylation of the phosphate group using trimethylsilyl diazomethane (TMS-diazomethane) improves chromatographic separation between BMP and PG, enabling improved distinction between BMP and PG isomers for LC/MS^38,39^. However, this method alters the original cellular lipidome and creates challenges for untargeted lipidomics, while also requiring additional time and experimental procedure. Instead, the most promising strategy for both distinguishing and quantifying BMP and PG has been liquid chromatography followed by tandem mass spectrometry (LC/MS2). Positive mode tandem mass spectrometry yields different fragmentation patterns for BMP and PG as ammonium adduct ions, and allows for clear distinction between the two. Since the first reporting of this distinction and technique, LC/MS2 has been used extensively with both untargeted and targeted mass spectrometry^16^. However, even with this MS2-based annotation technique, there are challenges to profiling BMPs that can continue to confound research if not carefully evaluated. Most prominently, the lack of individual standards for the many different BMP species makes it challenging to confirm the identity of most annotated BMPs.

Therefore, a robust approach is required to reliably distinguish and annotate BMPs. Here, we developed methods for systematically separating and relatively quantifying individual BMP and PG species using liquid chromatography/mass spectrometry (LC/MS) (Fig. 1). Our approach leverages both chemical and biological insights, which include: 1) differential molecular polarity, 2) tandem mass spectrometry (MS2) fragmentation patterns, 3) depletion of BMPs in cells lacking the recently discovered BMP synthase, ceroid lipofuscinosis 5 protein (CLN5)^40^, and 4) subcellular lipid enrichment patterns via rapid lysosome immunoprecipitation (LysoIP)^41^. Our method details experimental protocols for two state-of-the-art LC/MS instruments, the Orbitrap ID-X Tribrid and Ultivo triple quadruple (QQQ) mass spectrometers, and can be easily extended to other untargeted and targeted LC/MS instrumentation. Our work lays the foundations for accurately detecting and measuring BMPs in biological systems, facilitating studies in lysosomal and lipid biology, as well as disease and biomarker research.

**Figure 1.**
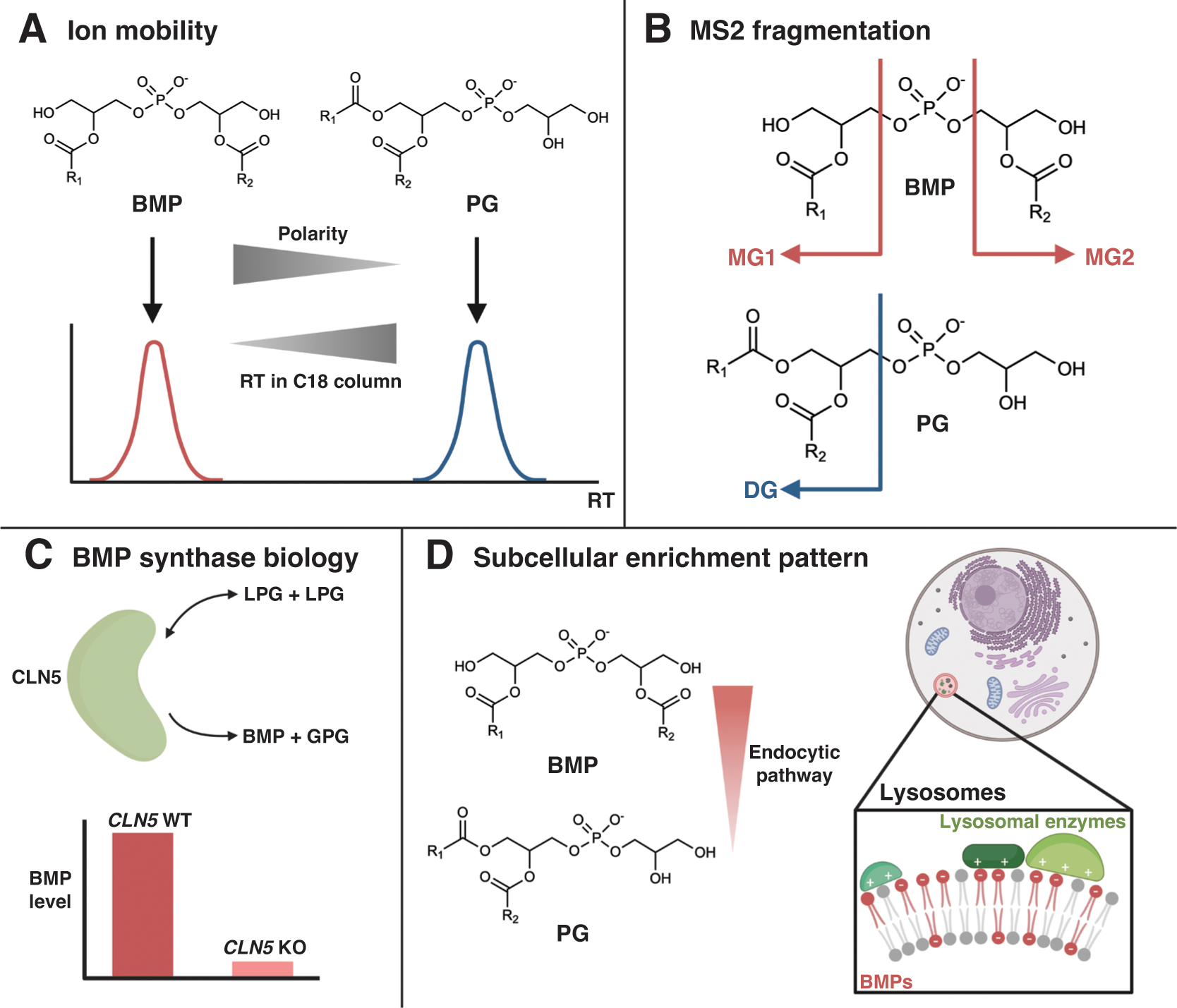
Principles for distinguishing BMPs and PGs. A. BMP and PG elute at different retention times (RTs) when utilizing an optimized chromatographic gradient, which is attributed to their isomeric differences in polarity. BMP is more polar than PG and thus elutes first as a distinct peak. B. BMP and PG exhibit different MS2 fragmentation patterns due to structural differences in acyl chain connectivity. BMP yields prominent MG fragments, while PG produces a prominent DG fragment. C. BMP and PG are synthesized by differing enzymes, with BMP’s recently discovered BMP synthase identified as CLN5. By knocking out CLN5, we can selectively deplete BMP and distinguish BMP species from PG species. D. BMP and PG have unique subcellular enrichment patterns, with BMP being distinctly enriched in lysosomes. By using LysoIP, we can identify BMP due to this characteristic lysosomal enrichment. MG = monoacylglycerol; DG = diacylglycerol; LPG = lysophosphatidylglycerol; GPG = glycerophosphoglycerol.

### Development of the protocol

This improved method for characterizing lysosomal BMP abundances was motivated by our overarching objective to characterize BMP levels during endocytic lipidomic reprogramming in neurodegeneration including neuronal ceroid lipofuscinoses (NCLs), CLN3 and CLN5 disease, in particular. The functions of the CLN3 and CLN5 proteins were unknown when we initiated our investigations, but we showed through our work that BMPs are reduced in CLN3-deficient lysosomes^30^, affirming previously reported results^31^. More importantly, when we moved on to our characterizations of CLN5, differentiating between BMP and PG was key to our discovery of CLN5 as the BMP synthase^40^. Strikingly, knocking out *CLN5* leads to markedly reduced BMP levels, at both the lysosomal and whole-cell levels. Importantly, this phenomenon has been observed not only using our method, but also by similar yet distinct patented methods in use by Nextcea, Inc. (Fig. S1), further strengthening the evidence of CLN5 as the BMP synthase.

During our initial untargeted lipidomic explorations of the function of CLN5, the discovery of changes in BMP levels was elusive since BMP can be difficult to distinguish from its isomeric lipid class PG. By manually scrutinizing differences in retention time (RT) and fragmentation patterns, we found that BMP and PG isomers can be distinguished with careful liquid chromatography design and MS2 analysis during positive mode fragmentation in the presence of ammonium on our Orbitrap ID-X Tribrid spectrometer. We further improved upon our methods for differentiating PG and BMP by performing targeted lipidomics on an Ultivo triple quadrupole (QQQ) mass spectrometer that targets characteristic fragments of BMPs and PGs. Using the Orbitrap and QQQ spectrometers, we were able to manually annotate and differentially analyze BMP and PG species in lysosomal and whole-cell tissue samples. However, we were still unable to conclusively differentiate between BMP and PG isomers for peaks that did not trigger MS2 fragmentation.

To overcome this challenge, we took advantage of CLN5 being the BMP synthase^40^ and that BMPs are enriched in lysosomes to formulate the BMP Identification Index (BMPII) and BMP Enrichment Score (BMPES), respectively. By doing so, we classified lipid targets that decreased substantially in the *CLN5* knockout (KO) relative to the wild-type (WT), yielding significantly low BMPII, and exhibited lysosomal enrichment, yielding high BMPES, as BMPs. These metrics, in conjunction with the detailed protocols for LC/MS analysis from both the Orbitrap and QQQ spectrometers, form a robust workflow for studying a diverse array of samples, spanning from lysosomal and cellular extracts from cell culture and animal tissues to patient-derived materials.

### Overview of the Protocol

This protocol comprises four major parts: 1) sample harvesting for whole-cells/tissue with optional LysoIP, 2) lipid extraction and LC/MS sample preparation, 3) Orbitrap setup and data analysis, and 4) QQQ setup and data analysis. This protocol establishes LC/MS methods for both an Orbitrap spectrometer and a QQQ spectrometer, while providing distinct MS2 spectra and quantitative parameters from both instruments, respectively. This protocol distinguishes itself from previous work through the development of the BMP identification index (BMPII) by perturbing the BMP synthase, as well as the BMP enrichment score (BMPES) by selectively enriching lysosomal BMP. These metrics act as valuable tools for definitively annotating and differentiating BMP and PG species, further distinguishing this protocol due its integration of biological insights to inform the analytical techniques used.

### Applications and Limitations

Our method relies on both the chemistry and biology of BMP to reliably identify BMPs using LC/MS. By optimizing our LC gradient, we allow for clean separation between BMP and PG, creating distinct peaks at different retention times (RT). We also use positive ion MS2 for characteristic BMP and PG fragmentation. BMP annotation can also be validated by calculating the change of candidate peak area in lipid standards extracted from BMP synthase (*CLN5*) KO against WT cells. Furthermore, by understanding the biology of BMP localization and synthesis, we can identify BMPs through their enrichment in lysosomal fractions via LysoIP.

While this method was first developed for cell culture samples, we have also adapted it to be used for animal tissue samples and patient-derived samples. As we continue to use and develop our BMP identification and quantitation method, we hope to expand its sample type coverage. An important limiting factor in the profiling of BMP is its low whole-cell abundance. Our method uses LysoIP as an optional tool to mitigate this issue. However, the LysoIP protocol requires prior technical skill and time. Thus, we primarily propose the use of premade BMPII standards of *CLN5* KO and WT, which can be manually created or acquired from others, to run on LC/MS alongside samples.

Additionally, the common limitations of LC/MS apply to our method as well. In particular, the expenses required for acquiring and maintaining mass spectrometry instrumentation may be a hurdle for some researchers. Availability of LC/MS at research institutions may also be an important limiting consideration, especially given the instrument time needed for proper LC separation of BMP and PG species. Potential avenues for improving throughput may explore shotgun mass spectrometry or expediting LC time for other lipid species if they are not targets of interest.

Finally, though our method is well established to distinguish BMPs from PGs in all acyl chain combinations, the method cannot differentiate regioisomers and stereoisomers of a BMP species. Thus, the method yields the total BMP level of a BMP species, resulting from the summation of BMP regioisomeric and stereoisomeric permutations. Current and future expansion of the technique is exploring this ability to distinguish BMP regioisomers and stereoisomers in order to investigate BMP eutomers and distomers in relation to specific disease-relevant biological activities.

### Materials

#### Reagents

Cells or animal tissue samples. Regular whole-cell or whole-tissue samples can be obtained from a variety of biological systems of interest.

**CAUTION** When working with biological samples, including animals and tissue samples, make sure to follow all applicable ethics and safety guidelines and regulations.

BMPII standards. Specifically, prepare or acquire lipidomics samples from *CLN5*

KO and WT cells ready for LC/MS.

**CAUTION** When working with biological samples, including animals and tissue samples, make sure to follow all applicable ethics and safety guidelines and regulations.

0.9% NaCl (w/v) saline solution (VWR, cat. no. S5815)
Optima LC/MS water (Fisher, cat. no. W6-4)
Optima LC/MS acetonitrile (Fisher, cat. no. A955-4)

**CAUTION** Acetonitrile is toxic. Wear proper personal protective equipment and follow all applicable chemical safety procedures when handling acetonitrile.

Optima LC/MS 2-propanol (IPA) (Fisher, cat. no. A461-500)

**CAUTION** 2-propanol is toxic. Wear proper personal protective equipment and follow all applicable chemical safety procedures when handling 2-propanol.

Optima LC/MS methanol (Fisher, cat. no. A456-4)

**CAUTION** Methanol is toxic. Wear proper personal protective equipment and follow all applicable chemical safety procedures when handling methanol.

Chloroform (Fisher, cat. no. C606SK-4)

**CAUTION** Chloroform is toxic. Wear proper personal protective equipment and follow all applicable chemical safety procedures when handling chloroform.

Ammonium formate (Millipore Sigma, cat. no. 70221-100G-F)
Formic acid (Fisher, cat. no. A117-50)
SPLASH LIPIDOMIX internal standard mix (Avanti, cat. no. 330707-1EA) **CAUTION** SPLASH LIPIDOMIX internal standard mix is a solution of lipids in methanol. Methanol is toxic. Wear proper personal protective equipment and follow all applicable chemical safety procedures when handling methanol.
[For tissue harvesting] Phosphate buffered saline, PBS (Thermo Fisher Scientific, cat. no. 10010023)
[For tissue harvesting or optional LysoIP protocol] Milli-Q water
[For tissue harvesting or optional LysoIP protocol] Deionized (DI) water
[For optional LysoIP protocol] Anti-HA magnetic beads (Thermo Fisher Scientific, cat. no. 88837)
[For optional LysoIP protocol] Potassium chloride, KCl (Sigma-Aldrich, cat. no. P9333-500G)
[For optional LysoIP protocol] Potassium phosphate, KH_2_PO_4_ (Millipore Sigma, cat. no. 57618)
[For optional LysoIP protocol] Potassium hydroxide, KOH (Fisher, cat. no. AC437131000)

#### Equipment

Benchtop centrifuge
[For adherent cell harvesting] Cell scrapers (Thermo Fisher Scientific, cat. no. 179707PK)
Micropipettes
Pipette tips
Microcentrifuge tube rack
1.5 mL tubes
SpeedVac vacuum concentrator (Labconco, cat. no. 7315060)
-80 °C Freezer
Sonicator
LC/MS autosampler caps (Fisher, cat. no. 6ASC9ST1x) LC/MS 2 mL autosampler glass vials (Fisher, cat. no. 6PSV9-1Px) and LC/MS 2 mL autosampler glass inserts (Fisher, cat. no. 6PME03C1SPx), or LC/MS 0.3 mL autosampler glass microvials (Fisher, cat. no 6PSV9-03FIVx)
Orbitrap ID-X Tribrid mass spectrometer (Thermo Fisher Scientific) equipped with a heated electrospray ionization (HESI) probe (RRID:SCR_025712)
Vanquish HPLC (Thermo Fisher Scientific) (RRID:n/a)
Ascentis Express C18 150 x 2.1 mm column (Millipore Sigma, cat. no. 53825-U) (RRID:n/a)
C18 5 x 2.1 mm guard (Sigma-Aldrich, cat. no. 53500-U) and cartridge (Sigma-Aldrich, cat. no. 50542-U) (RRID:n/a)
Agilent UHPLC guard, Eclipse Plus C18, 2.1mm, 1.8µm (Agilent Technologies 821725-901) (RRID:n/a)
Ultivo Triple Quadrupole mass spectrometer (Agilent Technologies) equipped with electrospray ionization (ESI) probe (Agilent G6465B) (RRID:n/a)
1290 Infinity II HPLC system (Agilent Technologies) including 1290 High Speed Pump (Agilent G7120A), 1290 Multisampler (Agilent G7167B), and 1290 MCT (G7116B) (RRID:SCR_019375)
[For tissue harvesting or optional LysoIP protocol] Glass vessel (VWR, cat no. 89026-386)
[For tissue harvesting or optional LysoIP protocol] Dounce homogenizer (douncer) (VWR, cat no. 89026-398)
[For optional LysoIP protocol] DynaMag spin magnet (magnet holder) (Thermo Fisher Scientific, cat. no. 12320D)
[For optional LysoIP protocol] Cell lifters (Corning, cat. no. 3008)
[For optional LysoIP protocol] Laboratory rocker
[For optional LysoIP protocol] 2 mL tubes

#### Software

LipidSearch (Thermo Fisher Scientific, cat. no. OPTON-30880) (RRID:n/a)
TraceFinder (Thermo Fisher Scientific, cat. no. OPTON-31001) (RRID:SCR_023045)
[Optional] FreeStyle (Thermo Fisher Scientific, cat. no. XCALI-97994) (RRID:SCR_022877). Optionally use for additional Orbitrap data visualization.
MassHunter Qualitative Analysis (Agilent Technologies) (RRID:SCR_019081)
MassHunter QQQ Quantitative Analysis (Quant My-Way) (Agilent Technologies) (RRID:SCR_015040)

### Reagent setup

#### Lipidomic Mobile Phase A (MPA) for LC

MPA is 10mM ammonium formate and 0.1% formic acid dissolved in 60% LC/MS grade water and 40% LC/MS grade acetonitrile. A 2-liter bottle of this solution can be made by adding 1.26 g of ammonium formate, 1200 mL of LC/MS grade water, 800 mL of LC/MS grade acetonitrile, and 2 mL of formic acid.

**CAUTION** Chloroform is toxic. Wear proper personal protective equipment and follow all applicable chemical safety procedures when handling chloroform.

#### Lipidomic Mobile Phase B (MPB) for LC

MPB is 10mM ammonium formate and 0.1% formic acid dissolved in 90% LC/MS grade 2-propanol and 10% LC/MS grade acetonitrile. A 2-liter bottle of this solution can be made by adding 1.26 g of ammonium formate, 200 mL of LC/MS grade acetonitrile, 1800 mL of LC/MS grade 2-propanol, and 2 mL of formic acid. This buffer needs to be sonicated for 2-3 hours, until the ammonium formate is fully dissolved.

**CAUTION** 2-propanol and acetonitrile are toxic. Wear proper personal protective equipment and follow all applicable chemical safety procedures when handling 2- propanol and acetonitrile.

### 80% methanol for cell harvesting

To make a 1 L bottle of this solution, add 800 mL of LC/MS grade methanol and 200 mL of LC/MS grade water.

**CAUTION** Methanol is toxic. Wear proper personal protective equipment and follow all applicable chemical safety procedures when handling methanol.

#### 2:1 chloroform:methanol solution (v/v) for lipid extraction

To make a 900 mL bottle of this solution, add 600 mL of LC/MS grade chloroform and 300 mL of LC/MS grade methanol. Add 900 μL of the SPLASH LIPIDOMIX to obtain 750 ng/mL concentration of the internal standard mix.

**CAUTION** Chloroform and methanol are toxic. Wear proper personal protective equipment and follow all applicable chemical safety procedures when handling chloroform and methanol.

#### 13:6:1 ACN:IPA:H2O (v/v/v) final lipidomic buffer for reconstitution of dry lipids

To make a 1 L bottle of this solution, add 650 mL of the LC/MS grade acetonitrile, 300 mL of the LC/MS grade 2-propanol, and 50 mL of the LC/MS grade water.

**CAUTION** Acetonitrile and 2-propanol are toxic. Wear proper personal protective equipment and follow all applicable chemical safety procedures when handling 2- propanol and acetonitrile.

### [For optional LysoIP protocol] LC/MS grade KPBS for washing steps during LysoIP

This KPBS solution consists of 136 mM KCl and 10 mM KH_2_PO_4_ in Optima LC/MS water at a pH of 7.25. The pH of this solution should be adjusted using KOH. Always keep this buffer on ice to maintain a cold temperature throughout the protocol. For tissue harvesting, KPBS can be substituted with PBS. KPBS is necessary for the optional LysoIP protocol.

### Orbitrap equipment setup for untargeted lipidomics

Lipid profiling involves chromatographic separation of the lipids in a sample followed by mass spectrometry analysis for identification and quantitation. These individual processes were optimized for the characterization of BMP and its distinction from the structural isomer PG by tandem mass spectrometry. The Vanquish HPLC utilized for liquid chromatography and Orbitrap ID-X for mass spectrometry were connected as one integrated LC/MS system. The methods for setting up and conducting an unbiased differential profiling as described here were adapted from our previous work^42,43^.

### Chromatographic gradient for untargeted lipidomics

A Vanquish HPLC was used to separate lipids based on polarity. An Ascentis Express C18 150 x 2.1 mm column (Millipore Sigma 53825-U) coupled with a 5 x 2.1 mm guard assembly (Sigma-Aldrich 53500-U and 50542-U) was mounted on the instrument. The elution for the samples was conducted at a flow rate of 0.26 mL/min with a linear change in gradient as described below for a total duration of 40 min.

| Time Interval (min) | Gradient (% B) |
| --- | --- |
| 0-1.5 | 32 |
| 1.5-4 | 45 |
| 4-5 | 52 |
| 5-8 | 58 |
| 8-11 | 66 |
| 11-14 | 70 |
| 14-18 | 75 |
| 18-21 | 97 |
| 21-35 | 97 |
| 35-35.1 | 32 |
| 35.1-40 | 32 |

### Parameters for tandem mass spectrometry

An Orbitrap ID-X Tribrid mass spectrometer (Thermo Fisher Scientific) with heated-electrospray ionization was used according to the specified parameters below. An MS1 scan was done to obtain precursor ions and for lipid quantitation and data-dependent (dd) MS2 for lipid identification based on fragmentation patterns.

| Parameter | Value |
| --- | --- |
| Ion transfer tube temperature | 300 °C |
| Vaporizer temperature | 375 °C |
| RF lens | 0.4 U |
| Positive ion voltage | 3,250 V |
| Negative ion voltage | 3,000 V |
| Sheath gas | 40 U |
| Aux gas | 10 U |
| Sweep gas | 1 U |
| **MS1** |  |
| Microscans | 1 U |
| Orbitrap resolution | 120,000 |
| Automatic Gain Control (AGC) target | $4 \times 10^5$ |
| Maximum injection time | 50 ms |
| Scan range | 250-1500 *m/z* |
| **MS2** |  |
| Microscans | 1 U |
| Detector type | Orbitrap |
| Orbitrap resolution | 30,000 |
| AGC target | $5 \times 10^5$ |
| Maximum injection time | 54 ms |
| Cycle time | 1.5 s |
| Isolation window | 1 *m/z* |
| Activation type | Higher-energy collision dissociation (HCD) |
| HCD fragmentation (%) | Stepped 15, 25, 35 |
| Intensity threshold | $1 \times 10^4$ |
| **dd settings** |  |
| Isotope exclusion | On |
| Dynamic exclusion time | 2.5 s |

### QQQ equipment setup for targeted lipidomics

Similar to the untargeted Orbitrap method, targeted QQQ characterization of BMP and distinction from its structural isomer PG were optimized using chromatographic separation and QQQ mass spectrometry. The 1290 Infinity II HPLC System utilized for liquid chromatography and QQQ for mass spectrometry were connected as one integrated LC/MS system. The methods for setting up and conducting a targeted differential profiling as described here were adapted from our previous work^42,44^.

### Chromatographic gradient for targeted lipidomics

A 1290 Infinity II HPLC was used to separate lipids based on polarity. An Agilent RRHD Eclipse Plus C18, 2.1mm, 100mm, 1.8µm column (Agilent Technologies 821725-901) coupled with a C18, 2.1mm, 1.8µm guard (Agilent Technologies 821725-901) was mounted on the instrument. The elution for the samples was conducted at a flow rate of 0.4 mL/min with a linear change in gradient as described below for a total duration of 16 minutes plus 2 minutes post time.

| Time Interval (min) | Gradient (% B) |
| --- | --- |
| 0-3 | 15 |
| 3-14 | 70 |
| 14-15 | 100 |
| 15-15.2 | 100 |
| 15.2-16 | 15 |
| Post time | 15 |

### Parameters for QQQ mass spectrometry LC/MS2

An Ultivo QQQ mass spectrometer (Agilent Technologies) with AJS-electrospray ionization (ESI) was used according to the specified parameters below. Lipids were identified using optimized multiple reaction monitoring (MRM) based on fragmentation patterns summarized under the MRM table. The MRMs included in the method were selected as a subset from those listed in the MRM table below.

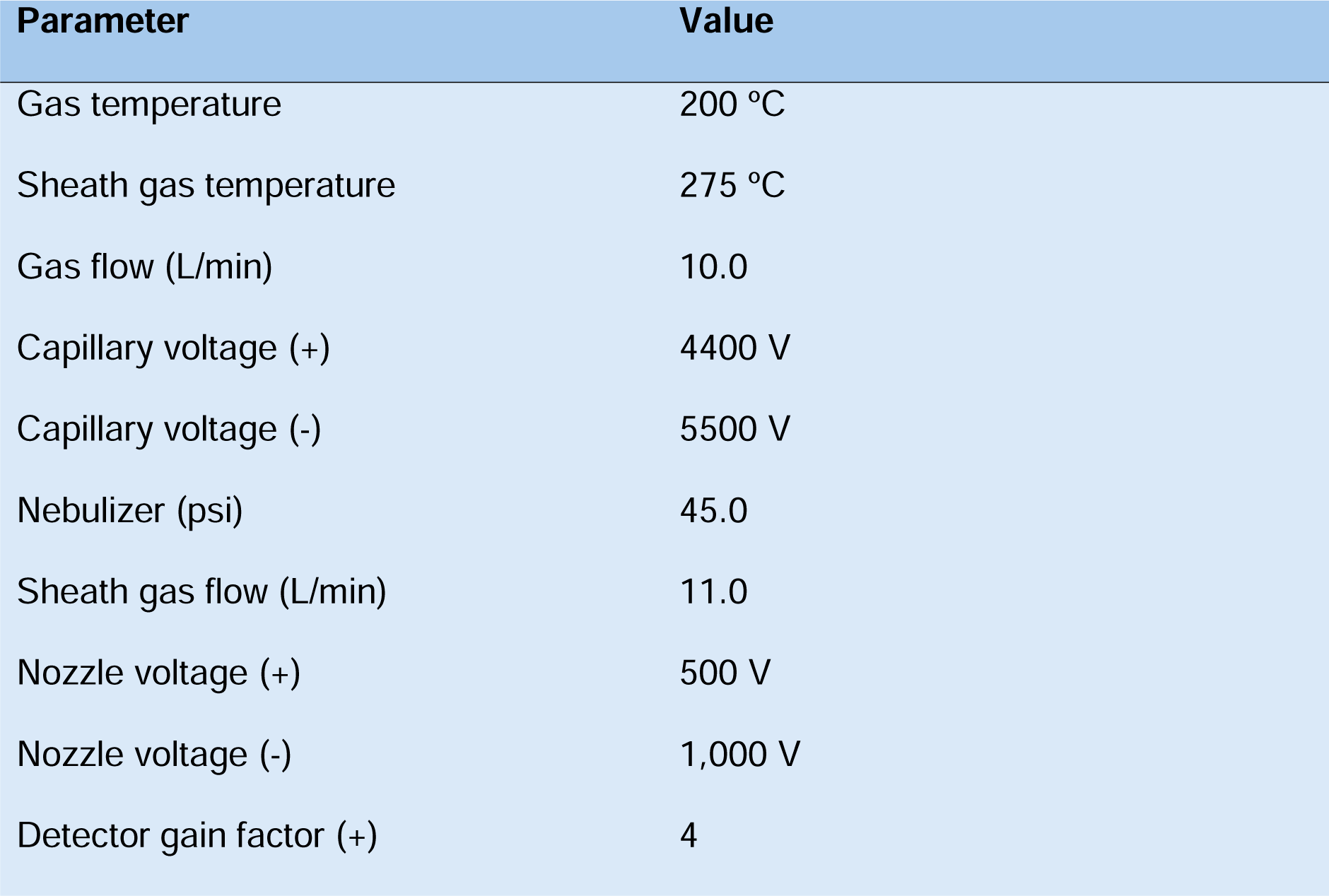
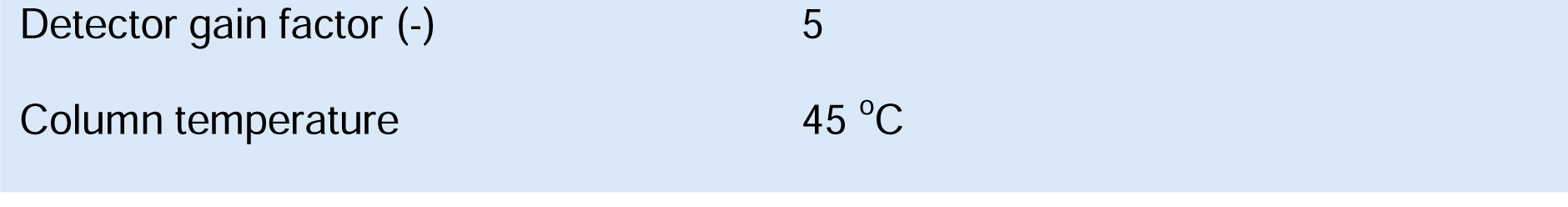

### Multiple Reaction Monitoring (MRM) transitions

| **Lipid** | **Precursor**<br>**Ion m/z**<br>**[M+NH<sub>4</sub>]<sup>+</sup>** | **Product**<br>**Ion m/z** | **Fragmentor**<br>**Voltage (V)** | **Collision**<br>**Energy (V)** | **Polarity** |
| --- | --- | --- | --- | --- | --- |
| BMP(14:0_16:0) | 712.5 | 285.2 | 150 | 27 | Positive |
| BMP(14:0_16:0) | 712.5 | 313.3 | 150 | 27 | Positive |
| BMP(14:0_18:1) | 738.5 | 285.2 | 150 | 27 | Positive |
| BMP(14:0_18:1) | 738.5 | 339.3 | 150 | 27 | Positive |
| BMP(16:0_16:0) | 740.5 | 313.3 | 150 | 27 | Positive |
| BMP(16:0_16:1) | 738.5 | 313.3 | 150 | 27 | Positive |
| BMP(16:0_16:1) | 738.5 | 311.3 | 150 | 27 | Positive |
| BMP(16:0_18:0) | 768.6 | 313.3 | 150 | 27 | Positive |
| BMP(16:0_18:0) | 768.6 | 341.3 | 150 | 27 | Positive |
| BMP(16:0_18:1) | 766.6 | 313.3 | 150 | 27 | Positive |
| BMP(16:0_18:1) | 766.6 | 339.3 | 150 | 27 | Positive |
| BMP(16:0_20:4) | 788.5 | 313.3 | 150 | 27 | Positive |
| BMP(16:0_20:4) | 788.5 | 361.3 | 150 | 27 | Positive |
| BMP(16:0_22:6) | 812.5 | 313.3 | 150 | 27 | Positive |
| BMP(16:0_22:6) | 812.5 | 385.3 | 150 | 27 | Positive |
| BMP(16:1_16:1) | 736.5 | 311.3 | 150 | 27 | Positive |
| BMP(16:1_18:1) | 764.5 | 311.3 | 150 | 27 | Positive |
| BMP(16:1_18:1) | 764.5 | 339.3 | 150 | 27 | Positive |
| BMP(16:1_18:2) | 762.5 | 311.3 | 150 | 27 | Positive |
| BMP(16:1_18:2) | 762.5 | 337.3 | 150 | 27 | Positive |
| BMP(16:1_20:3) | 788.5 | 311.3 | 150 | 27 | Positive |
| BMP(16:1_20:3) | 788.5 | 363.3 | 150 | 27 | Positive |
| BMP(16:1_22:6) | 810.5 | 311.3 | 150 | 27 | Positive |
| BMP(16:1_22:6) | 810.5 | 385.3 | 150 | 27 | Positive |
| BMP(18:0_18:0) | 796.6 | 341.3 | 150 | 27 | Positive |
| BMP(18:0_18:1) | 794.6 | 341.3 | 150 | 27 | Positive |
| BMP(18:0_18:1) | 794.6 | 339.3 | 150 | 27 | Positive |
| BMP(18:1_18:1) | 792.6 | 339.3 | 150 | 27 | Positive |
| BMP(18:1_18:2) | 790.6 | 339.3 | 150 | 27 | Positive |
| BMP(18:1_18:2) | 790.6 | 337.3 | 150 | 27 | Positive |
| BMP(18:1_20:0) | 822.6 | 339.3 | 150 | 27 | Positive |
| BMP(18:1_20:0) | 822.6 | 369.3 | 150 | 27 | Positive |
| BMP(18:1_20:1) | 820.6 | 339.3 | 150 | 27 | Positive |
| BMP(18:1_20:1) | 820.6 | 367.3 | 150 | 27 | Positive |
| BMP(18:1_20:2) | 818.6 | 339.3 | 150 | 27 | Positive |
| BMP(18:1_20:3) | 816.6 | 363.3 | 150 | 27 | Positive |
| BMP(18:1_20:4) | 814.6 | 339.3 | 150 | 27 | Positive |
| BMP(18:1_20:4) | 814.6 | 361.3 | 150 | 27 | Positive |
| BMP(18:1_22:1) | 848.6 | 339.3 | 150 | 27 | Positive |
| BMP(18:1_22:1) | 848.6 | 395.4 | 150 | 27 | Positive |
| BMP(18:1_22:2) | 846.6 | 339.3 | 150 | 27 | Positive |
| BMP(18:1_22:2) | 846.6 | 393.3 | 150 | 27 | Positive |
| BMP(18:1_22:3) | 844.6 | 339.3 | 150 | 27 | Positive |
| BMP(18:1_22:3) | 844.6 | 391.3 | 150 | 27 | Positive |
| BMP(18:1_22:4) | 842.6 | 339.3 | 150 | 27 | Positive |
| BMP(18:1_22:4) | 842.6 | 389.3 | 150 | 27 | Positive |
| BMP(18:1_22:5) | 840.6 | 339.3 | 150 | 27 | Positive |
| BMP(18:1_22:5) | 840.6 | 387.3 | 150 | 27 | Positive |
| BMP(18:1_22:6) | 838.6 | 339.3 | 150 | 27 | Positive |
| BMP(18:1_22:6) | 838.6 | 385.3 | 150 | 27 | Positive |
| BMP(18:2_18:2) | 788.5 | 337.3 | 150 | 27 | Positive |
| BMP(18:2_20:3) | 814.6 | 337.3 | 150 | 27 | Positive |
| BMP(18:2_20:3) | 814.6 | 363.3 | 150 | 27 | Positive |
| BMP(18:2_22:6) | 836.5 | 337.3 | 150 | 27 | Positive |
| BMP(18:2_22:6) | 836.5 | 385.3 | 150 | 27 | Positive |
| BMP(20:4_20:4) | 836.5 | 361.3 | 150 | 27 | Positive |
| BMP(20:4_22:6) | 860.5 | 361.3 | 150 | 27 | Positive |
| BMP(20:4_22:6) | 860.5 | 385.3 | 150 | 27 | Positive |
| BMP(20:5_22:6) | 858.5 | 359.3 | 150 | 27 | Positive |
| BMP(20:5_22:6) | 858.5 | 385.3 | 150 | 27 | Positive |
| BMP(22:4_22:6) | 888.6 | 389.3 | 150 | 27 | Positive |
| BMP(22:4_22:6) | 888.6 | 385.3 | 150 | 27 | Positive |
| BMP(22:5_22:5) | 888.6 | 387.3 | 150 | 27 | Positive |
| BMP(22:5_22:6) | 886.6 | 387.3 | 150 | 27 | Positive |
| BMP(22:5_22:6) | 886.6 | 385.3 | 150 | 27 | Positive |
| BMP(22:6_22:6) | 884.5 | 385.3 | 150 | 27 | Positive |
| PG(14:0_16:0) | 712.5 | 523.5 | 110 | 9 | Positive |
| PG(14:0_18:1;<br>16:0_16:1) | 738.6 | 549.5 | 110 | 9 | Positive |
| PG(16:0_16:0) | 740.5 | 551.5 | 110 | 9 | Positive |
| PG(16:0_18:0) | 768.6 | 579.5 | 110 | 9 | Positive |
| PG(16:0_18:1) | 766.6 | 577.5 | 110 | 9 | Positive |
| PG(16:0_20:4;<br>16:1_20:3;<br>18:2_18:2) | 788.5 | 599.5 | 110 | 9 | Positive |
| PG(16:0_22:6) | 812.5 | 623.5 | 110 | 9 | Positive |
| PG(16:1_16:1) | 736.5 | 547.5 | 110 | 9 | Positive |
| PG(16:1_18:1) | 764.5 | 575.5 | 110 | 9 | Positive |
| PG(16:1_18:2) | 762.5 | 573.5 | 110 | 9 | Positive |
| PG(16:1_22:6) | 810.5 | 621.5 | 110 | 9 | Positive |
| PG(18:0_18:0) | 796.6 | 607.6 | 110 | 9 | Positive |
| PG(18:0_18:1) | 794.6 | 605.6 | 110 | 9 | Positive |
| PG(18:1_18:1) | 792.6 | 603.5 | 110 | 9 | Positive |
| PG(18:1_18:2) | 790.6 | 601.5 | 110 | 9 | Positive |
| PG(18:1_20:0) | 822.6 | 633.6 | 110 | 9 | Positive |
| PG(18:1_20:1) | 820.6 | 631.6 | 110 | 9 | Positive |
| PG(18:1_20:2) | 818.6 | 629.6 | 110 | 9 | Positive |
| PG(18:1_20:3) | 816.6 | 627.5 | 110 | 9 | Positive |
| PG(18:1_20:4;<br>18:2_20:3) | 814.6 | 625.5 | 110 | 9 | Positive |
| PG(18:1_22:1) | 848.6 | 659.6 | 110 | 9 | Positive |
| PG(18:1_22:2) | 846.6 | 657.6 | 110 | 9 | Positive |
| PG(18:1_22:3) | 844.6 | 655.6 | 110 | 9 | Positive |
| PG(18:1_22:4) | 842.6 | 653.6 | 110 | 9 | Positive |
| PG(18:1_22:5) | 840.6 | 651.5 | 110 | 9 | Positive |
| PG(18:1_22:6) | 838.6 | 649.5 | 110 | 9 | Positive |
| PG(18:2_22:6;<br>20:4_20:4) | 836.5 | 647.5 | 110 | 9 | Positive |
| PG(20:4_22:6) | 860.5 | 671.5 | 110 | 9 | Positive |
| PG(20:5_22:6) | 858.5 | 669.5 | 110 | 9 | Positive |
| PG(22:4_22:6;<br>22:5_22:5) | 888.6 | 699.5 | 110 | 9 | Positive |
| PG(22:5_22:6) | 886.6 | 697.5 | 110 | 9 | Positive |
| PG(22:6_22:6) | 884.5 | 695.5 | 110 | 9 | Positive |
| PG(15:0_18:1-d7) | 759.6 | 570.5 | 110 | 9 | Positive |
| PC(16:0_18:1) | 760.6 | 86.1 | 215 | 50 | Positive |
| PC(16:0_18:1) | 760.6 | 184.1 | 215 | 50 | Positive |
| PC(16:0_18:1) | 760.6 | 478.3 | 215 | 50 | Positive |
| PC(16:0_18:1) | 760.6 | 504.3 | 215 | 50 | Positive |
| PC(18:1_18:1) | 786.6 | 86.1 | 215 | 50 | Positive |
| PC(18:1_18:1) | 786.6 | 184.1 | 215 | 50 | Positive |
| PC(18:1_18:1) | 786.6 | 504.3 | 215 | 50 | Positive |
| PC(15:0_18:1-d7) | 753.6 | 86.1 | 215 | 50 | Positive |
| PC(15:0_18:1-d7) | 753.6 | 184.1 | 215 | 50 | Positive |
| PC(15:0_18:1-d7) | 753.6 | 464.4 | 215 | 50 | Positive |
| PC(15:0_18:1-d7) | 753.6 | 511.4 | 215 | 50 | Positive |

#### Procedure

Sample harvesting for lipidomic analysis

**TIMING** 4 h

1. Different procedures are detailed below depending on sample types. Follow option A for BMP analysis from adherent cell culture. Follow option B for BMP analysis from mouse tissues. For extra validation through LysoIP, follow option C for LysoIP with adherent cell culture or option D for LysoIP with mouse tissues.

**CRITICAL** Prior to harvesting, plan for sample normalization by growing additional wells for BCA protein assay, cell count, or other equivalent measurements.

(A) Harvesting of whole-cell samples for BMP analysis

The procedure in 1A is optimized specifically for harvesting whole-cell lipids from adherent cells in 6-well plates. To harvest cells from other well plates or dishes, scale the volumes of solvents accordingly as needed.

i. Pre-chill a benchtop centrifuge to 4 °C.
ii. Retrieve both dry ice and ice in buckets.
iii. Set cell scrapers ready for fast harvesting of lipid samples.
iv. At the time of harvesting, place plates of cells on ice and aspirate media.

**CAUTION** When working with biological samples, make sure to follow all applicable ethics and safety guidelines and regulations.

Wash each well one time with 1.5 mL of ice-cold 0.9% NaCl (w/v) and aspirate.
Transfer plates to dry ice.
Add 100 μL of 80% methanol to each well.

**CAUTION** Methanol is toxic. Wear proper personal protective equipment and follow all applicable chemical safety procedures when handling methanol.

(viii) For each plate, hold the plate at a slight angle and use a pipette tip attached to a micropipette to thoroughly scrape each well. Ensure the tip is held perpendicular to prevent it from falling off the pipette. Then, use a cell scraper to thoroughly collect the cells at the bottom of each well. **CRITICAL STEP** This step must be performed consistently across all wells. It is recommended to follow a specific pattern of scraping across all wells to minimize variance.
(ix) Transfer the scraped cell harvest of each well into its own respective pre-chilled 1.5 mL tube with a micropipette. These tubes will be used for lipid extraction.

**PAUSE POINT** Cell harvests are stable for storage and can be stored at -80 °C.

(B) Harvesting of mouse whole-tissue samples for BMP analysis

The procedure in 1B is optimized specifically for harvesting whole-tissue lipids from mouse brain or liver. To harvest other tissues from mice or tissues from different model organisms, scale volumes as needed.

**CAUTION** When working with animals and tissue samples, make sure to follow all applicable ethics and safety guidelines and regulations.

i. Pre-chill a benchtop centrifuge to 4 °C.
ii. Wash the douncer that will be used to homogenize and lyse the cells.

Wash each glass vessel 10 times with deionized (DI) and Milli-Q water. Leave them drying upside down on a paper towel and then pre-chill the glass vessel on ice.

**CRITICAL** Always avoid any contact with the tissue grinding part of the douncer as it will be directly touching cells.

(iii) At the time of harvesting, add 950 μL of cold PBS to the glass vessel of the douncers and pre-chill.
(iv) Sacrifice animal(s) and dissect tissues of interest. For mouse brains, collect following euthanasia, dissect cerebral hemispheres on an ice-cold plastic dish, and use half cerebral hemisphere for each sample. For mouse livers, collect following euthanasia and isolate a small round piece of liver using a biopsy punch with a 4 mm diameter for each sample. **CAUTION** When working with animals and tissue samples, make sure to follow all applicable ethics and safety guidelines and regulations.
(v) Transfer tissues immediately after dissecting to the glass vessels of the douncers.
(vi) Gently dounce the tissues 25 times on ice, avoiding making bubbles as much as possible.

**CRITICAL STEP** This homogenizes the tissue suspension and mechanically breaks the plasma membrane. It is important to homogenize the same number of times and at a consistent speed across all samples.

(vii) Use a serological pipette to transfer all the homogenized samples from the glass vessels into new 1.5 mL tubes.
(viii) Transfer 100 μL of each sample into its own respective pre-chilled 1.5 mL tube with a micropipette. These tubes will be used for lipid extraction. **CRITICAL** Adjust the volume transferred for lipid extraction depending on initial LC/MS results. If lipid signal is too low, increase the volume transferred. If lipid signal is saturating, decrease the volume transferred. PAUSE POINT Tissue harvests are stable for storage and can be stored at -80 °C.

(C) Harvesting of lysosomes from cells for BMP analysis by LysoIP

The procedure in 1C is optimized specifically for harvesting lysosomal lipids from adherent HEK293T cells (RRID:CVCL_0063) expressing 3xHA-tagged TMEM192 protein in 15 cm plates. If using a different cell line, the LysoIP protocol needs to be optimized^45^. To harvest cells from other plates, well plates, or dishes, scale the volumes of solvents accordingly as needed.

**CRITICAL** Cells can be used for LysoIP if they express the 3xHA-tagged TMEM192 construct. Cell lines expressing 3xHA-tagged TMEM192 can be generated by lentiviral transduction of the pLJC5-Tmem192-3xHA plasmid construct (RRID:Addgene_102930)^41^.

i. Pre-chill a benchtop centrifuge to 4 °C.
ii. Wash the douncer that will be used to homogenize and lyse the cells.

Wash each glass vessel 10 times with DI and Milli-Q water. Leave them drying upside down on a paper towel and then pre-chill the glass vessel on ice.

**CRITICAL** Always avoid any contact with the tissue grinding part of the douncer as it will be directly touching cells.

(iii) Prepare one set of 6 microcentrifuge tubes for each cell plate to be harvested. For example, if 6 plates are harvested, 6 sets of tubes should be prepared as shown below for a total of 36 tubes. Set them on a tube rack on ice from left-to-right, as follows:

Tube #1: 2 mL tube for initial cell harvest.

Tube #2: 1.5 mL tube for homogenized suspension after douncing. Tube #3: 1.5 mL tube for whole-cell (WC) sample.

Tube #4: 1.5 mL tube for magnetic beads.

Tube #5: 1.5 mL tube for post-magnetic sample.

Tube #6: 1.5 mL tube for final LysoIP sample

(iv) Pipette the total required volume of anti-HA magnetic beads needed for the experiment into an appropriately sized tube. Each cell plate will require 100 μL of magnetic beads. For example, if 6 plates are harvested, 600 μL total should be pooled into a separate container. Be sure to shake the bottle of beads well before pipetting as beads tend to settle at the bottom of the container.
(v) Add an equivalent amount of cold KPBS to the magnetic beads. For example, if there is 600 μL of magnetic beads, add 600 μL of KPBS. Then pipette up and down one time.
(vi) Place the tube on the magnet holder. Aspirate any liquid from the bottom and sides of the tube not adjacent to the magnet, avoiding the beads.
(vii) Remove the tube from the magnet holder. Repeat the wash and aspiration steps 1C(v-vi) two times, for a total of three washes.
(viii) Remove the tube from the magnet holder. Resuspend the beads with an equivalent volume of KPBS, as described in step 1C(v), and aliquot 100 μL into each of the #4 tubes prepared earlier.
(ix) At the time of harvesting, move the first set of plates to the bench and set them on ice.

**CRITICAL** This protocol suggests preparing and harvesting two 15 cm plates at a time for both efficiency and quality. To achieve the best quality of LysoIP, harvest one plate at a time.

**CAUTION** When working with biological samples, make sure to follow all applicable ethics and safety guidelines and regulations.

Decant the media and wash the cells two times with ice-cold KPBS. For the first wash, gently pour ∼5 mL of the KPBS along the edges of the plate to decant it. For the second wash, aspirate the KPBS instead.
Add 950 μL of cold KPBS to each plate.
Hold the plate at a slight angle and use a cell lifter to scrape the cells to the bottom of the plate. Visually check that all cells have been harvested. **CRITICAL STEP** This step must be performed consistently across all wells. It is recommended to follow a specific pattern of scraping across all wells to minimize variance.
Transfer the detached cell suspension of each plate into its own respective tube #1 with a micropipette.
Centrifuge the cell suspension in tubes #1 at 1,000g and 4 °C for 2 minutes.
Aspirate the supernatant with a micropipette and resuspend the cells with 950 μL of cold KPBS. If using a pellet mixer for homogenization, first resuspend the cells with 100 μL of cold KPBS and homogenize. Then, fill the tube to 950 μL and proceed with the protocol.
Transfer 25 μL of the resuspended cells into tube #3 for whole-cell (WC) processing. The remaining 925 μL suspension will be used for generating the LysoIP samples.
Transfer the remaining 925 μL of cell suspension from tube #1 into the pre-chilled glass vessel of the douncer.
Gently dounce the tissues 25 times on ice, avoiding making bubbles.

**CRITICAL STEP** This step homogenizes the cell suspension and mechanically breaks the plasma membrane to release intracellular organelles including lysosomes. It is important to homogenize the same number of times and at a consistent speed across all samples.

(xix) Use a serological pipette to transfer the homogenized sample from the glass vessels into the respective tubes #2.
(xx) Centrifuge the tubes #2 homogenized suspensions at 1,000g and 4 °C for 2 minutes. While waiting for the centrifugation, wash the douncers for the

next set of harvestings. Take care to avoid physical contact with the tissue grinder part of the douncer that inserts into the glass vessel.

(xxi) Transfer the supernatant formed after centrifugation in tube #2 into each sample’s respective tube #4, which contains the magnetic beads. Pipette up and down one time to resuspend the mixture.

**CRITICAL** Organelles including lysosomes are contained in the supernatant after the spin-down. Be careful to avoid taking any insoluble material when transferring the supernatant, as this may affect the results.

(xxii) Rock tubes #4 gently in the cold room for 3 minutes.

**CRITICAL** Steps 1C(xxii-xxxii) should be carried out inside the cold room.

(xxiii) Place tubes #4 in the magnet holder and wait 25 seconds for the beads to be pulled by the magnet along the wall of the tube.

**CRITICAL** Maintain this timing consistently across all samples being harvested.

(xxiv) For the first wash, remove tube #4 from the magnet and add 1 mL of cold KPBS. Pipette up and down 2-3 times, consistent between washes. Next, place the tube back on the magnet and wait 25 seconds. Finally, aspirate any liquid from the bottom and sides of the tube not adjacent to the magnet, as well as any liquid that may have been trapped in the cap.
(xxv) For the second wash, remove tube #4 from the magnet and add 1 mL of cold KPBS. Pipette up and down 2-3 times, consistent between washes. Then, place the tube back on the magnet and wait 25 seconds. Finally,

aspirate any liquid from the bottom and sides of the tube not adjacent to the magnet.

(xxvi) For the third wash, remove tube #4 from the magnet and add 1 mL of cold KPBS. Pipette up and down 2-3 times, consistent with the washes. Then, transfer the resuspended solution to the respective tube #5.

**CRITICAL** Transferring the sample to empty tube #5 for the third and final wash gives a cleaner IP. The leftover beads in tube #5 after the final wash contain lysosomes.

(xxvii) Place tube #5 back on the magnet and wait 25 seconds. Finally, aspirate any liquid from the bottom and sides of the tube not adjacent to the magnet.
(xxviii) Remove tube #5 from the magnet. The leftover contents in tube #5 after this final wash contain only the beads and the lysosomes.
(xxix) Resuspend the bound beads and lysosomes in tube #5 with 1,000 μL of the 2:1 chloroform:methanol (v/v) solution containing internal standards (see “Reagent Setup”). This solution lyses the lysosome, allowing for extraction of the lipids present.

**CRITICAL STEP** Due to the difficulty of resuspending the bound beads, begin by flushing along the walls of the tube.

**CAUTION** Chloroform and methanol are toxic. Wear proper personal protective equipment and follow all applicable chemical safety procedures when handling chloroform and methanol.

(xxx) Leave the samples on ice and start the LysoIP process for the next set of plates.
(xxxi) After repeating steps 1C(ix-xxix) for all sets of plates: Tubes #3 should contain 25 μL of WC sample.

Tubes #5 should contain 1,000 μL of LysoIP sample (with beads still attached).

(xxxii) 10 minutes after the last LysoIP, place all tubes #5 in the magnet and wait 25 seconds. Then, transfer each tube’s supernatant to its corresponding tube #6, making sure to avoid the beads adjacent to the magnet. Tubes #6 now hold the final LysoIP samples, without the beads.

**PAUSE POINT** Cell harvests are stable for overnight storage at -80 °C.

**Harvesting of lysosomes from mouse tissues for BMP analysis by LysoIP** The procedure in 1D is optimized specifically for harvesting lysosomal lipids from brains or livers of mice expressing 3xHA-tagged TMEM192 protein. To harvest other tissues from mice or tissues from different model organisms, scale tissue volumes accordingly as needed.

**CRITICAL** Mouse tissues can be used for LysoIP if they express the 3xHA- tagged TMEM192 construct^30^.

**CAUTION** When working with animals and tissue samples, make sure to follow all applicable ethics and safety guidelines and regulations.

i. Pre-chill a benchtop centrifuge to 4 °C.
ii. Wash the douncer that will be used to homogenize and lyse the tissues.

Wash each glass vessel 10 times with DI and Milli-Q water. Leave them drying upside down on a paper towel and then pre-chill the glass vessel on ice.

**CRITICAL** Always avoid any contact with the tissue grinding part of the douncer as it will be directly touching cells.

(iii) Prepare one set of 6 microcentrifuge tubes for each tissue to be harvested. For example, if 6 tissues are harvested, 6 sets of tubes should be prepared as shown below for a total of 36 tubes. Set them on a tube rack on ice from left-to-right, as follows:

Tube #1: Not used in whole-tissue protocol but can be used for aliquoting whole-tissue samples from Tube #2.

Tube #2: 1.5 mL tube for homogenized suspension after douncing. Tube #3: 1.5 mL tube for whole-tissue sample.

Tube #4: 1.5 mL tube for magnetic beads.

Tube #5: 1.5 mL tube for post-magnetic sample. Tube #6: 1.5 mL tube for final LysoIP sample.

(iv) Pipette the total required volume of anti-HA magnetic beads needed for the experiment into an appropriately sized tube. Each tissue requires 100 μL of magnetic beads. For example, if 6 tissues are harvested, 600 μL total should be pooled into a separate container. Be sure to shake the bottle of beads well before pipetting as beads tend to settle at the bottom of the container.
(v) Add an equivalent amount of ice-cold KPBS to the magnetic beads. For example, if there is 600 μL of magnetic beads, add 600 μL of KPBS. Then pipette up and down one time.
(vi) Place the tube on the magnet holder. Aspirate any liquid from the bottom and sides of the tube not adjacent to the magnet, avoiding the beads.
(vii) Remove the tube from the magnet holder. Repeat the wash and aspiration steps 1D(v-vi) two times, for a total of three washes.
(viii) Remove the tube from the magnet holder. Resuspend the beads with an equivalent volume of KPBS, as described in step 1D(v), and aliquot 100 μL into each of the #4 tubes prepared earlier.
(ix) Resuspend the beads with an equivalent volume of KPBS, as described in the previous step, and aliquot 100 μL into each of the #4 tubes prepared earlier.
(x) At the time of harvesting, add 950 μL of cold KPBS to the glass vessel of the douncers and pre-chill.
(xi) Sacrifice animal(s) and dissect tissues of interest. For mouse brains, collect following euthanasia, dissect cerebral hemispheres on an ice-cold plastic dish, and use half cerebral hemisphere for each sample. For mouse livers, collect following euthanasia and isolate a small round piece of liver using a biopsy punch with a 4 mm diameter for each sample. **CAUTION** When working with animals and tissue samples, make sure to follow all applicable ethics and safety guidelines and regulations.
(xii) Transfer tissue immediately after dissecting to the glass vessels of the douncers.
(xiii) Gently dounce the tissues 25 times on ice, avoiding making bubbles.

**CRITICAL STEP** This step homogenizes the tissues and mechanically breaks the plasma membrane to release intracellular organelles including lysosomes. It is important to homogenize the same number of times and at a consistent speed across all samples.

(xiv) Use a serological pipette to transfer the homogenized sample from the glass vessels into the respective tubes #2.
(xv) Transfer 25 μL of the homogenized sample from each tube #2 to its own respective tube #3 for whole-tissue processing. The remaining 925 μL suspension will be used for generating the LysoIP samples.
(xvi) Centrifuge the tubes #2 homogenized suspensions at 1,000g and 4 °C for 2 minutes. While waiting for the centrifugation, wash the douncers for the next set of harvestings. Take care to avoid physical contact with the tissue grinder part of the douncer that inserts into the glass vessel.
(xvii) Transfer the supernatant formed after centrifugation in tube #2 into each sample’s respective tube #4, which contains the magnetic beads. Pipette up and down one time to resuspend the mixture.

**CRITICAL** Organelles including lysosomes are contained in the supernatant after the spin-down. Be careful to avoid taking any insoluble material when transferring the supernatant, as this may affect the results.

(xviii) Rock tubes #4 gently for 3 minutes in the cold room.

**CRITICAL** Steps 1D(xviii – xxviii) should be carried out inside the cold room.

(xix) Place tubes #4 in the magnet holder and wait 25 seconds for the beads to be pulled by the magnet along the wall of the tube.

**CRITICAL** Maintain this timing consistently across all samples being harvested.

(xx) For the first wash, remove tube #4 from the magnet and add 1 mL of cold KPBS. Pipette up and down 2-3 times, consistent between washes. Next, place the tube back on the magnet and wait 25 seconds. Finally, aspirate any liquid from the bottom and sides of the tube not adjacent to the magnet, as well as any liquid that may have been trapped in the cap.
(xxi) For the second wash, remove tube #4 from the magnet and add 1 mL of cold KPBS. Pipette up and down 2-3 times, consistent between washes. Then, place the tube back on the magnet and wait 25 seconds. Finally, aspirate any liquid from the bottom and sides of the tube not adjacent to the magnet.
(xxii) For the third wash, remove tube #4 from the magnet and add 1 mL of cold KPBS. Pipette up and down 2-3 times, consistent with the washes. Then, transfer the resuspended solution to the respective tube #5.

**CRITICAL STEP** Transferring the sample to empty tube #5 for the third and final wash gives a clearer result. The leftover beads in tube #5 after the final wash contain lysosomes.

(xxiii) Place the tube #5 back on the magnet and wait 25 seconds. Finally, aspirate any liquid from the bottom and sides of the tube not adjacent to the magnet.
(xxiv) Remove tube #5 from the magnet. The leftover contents in tube #5 after this final wash contain only the beads and the lysosomes.
(xxv) Resuspend the bound beads and lysosomes in tubes #5 with 1,000 μL of the 2:1 chloroform:methanol (v/v) solution containing internal standards (see “Reagent Setup”). This solution lyses the lysosome, allowing for extraction of the lipids present.

**CRITICAL STEP** Due to the difficulty of resuspending the bound beads, begin by flushing along the walls of the tube.

**CAUTION** Chloroform and methanol are toxic. Wear proper personal protective equipment and follow all applicable chemical safety procedures when handling chloroform and methanol.

(xxvi) Leave the samples on ice and start the LysoIP process for the next set of tissue samples.
(xxvii) After repeating steps 1D(x-xxv) for all tissues:

Tubes #3 should contain 25 μL of whole-tissue sample.

Tubes #5 should contain 1,000 μL of LysoIP sample (with beads still attached).

(xxviii) 10 minutes after the last LysoIP, place all tubes #5 in the magnet and wait 25 seconds. Then, transfer each tube’s supernatant to its corresponding

tube #6, making sure to avoid the beads adjacent to the magnet. Tubes #6 now hold the final LysoIP samples, without the beads.

**PAUSE POINT** Tissue harvests are stable for overnight storage at -80 °C.

#### Lipid extraction and processing from harvested samples

**TIMING** 4 h

**CRITICAL** The samples harvested thus far will be further processed for extraction before the LC/MS measurement. The specific buffers and processing steps to be followed vary depending on the polarity of the metabolites of interest. Since BMP is a glycerophospholipid, a lipidomic analysis is preferred here. The specific processing steps described in this section are optimized for lipid extraction and adapted according to our previous work^30,46^.

1. 2. Add 1,000 μL of 2:1 chloroform:methanol (v/v) solution containing internal standards (see “Reagent Setup”) to all whole-cell/whole-tissue tubes from 1A or 1B. For 1C and 1D, this specifically refers to tubes #3.

**CAUTION** Chloroform and methanol are toxic. Wear proper personal protective equipment and follow all applicable chemical safety procedures when handling chloroform and methanol.

1. 3. Vortex all samples at 4 °C for 1 hour. This includes both WC/whole-tissue and LysoIP samples. For the procedure with LysoIP, the LysoIP samples are tubes #6.
2. 4. Add 200 μL of 0.9% NaCl (w/v) solution to all samples.
3. 5. Vortex all samples at 4 °C for 10 minutes.
4. 6. Centrifuge all samples at 3,000g and 4 °C for 5 minutes.

**CRITICAL STEP** Two layers should be visible in these tubes after centrifugation: a polar layer at the top and a non-polar layer at the bottom containing the lipids.

Prepare a new set of 1.5 mL microcentrifuge tubes during this centrifugation for collecting the non-polar layer.

1. 7. Carefully collect the bottom layer containing the lipids and transfer it to the newly created set of 1.5 mL microcentrifuge tubes.
2. 8. Dry these samples in a SpeedVac vacuum concentrator for as long as needed for all liquid to evaporate, leaving the dried lipid pellets at the bottom of the tubes.

**PAUSE POINT** Dried lipids are the most stable for storage and can be stored at -80 °C.

1. 9. Reconstitute the dried lipids in the samples with 50 μL of the 13:6:1 ACN:IPA:H2O (v/v/v) lipidomic buffer.

**CRITICAL** Steps 9-14 should ideally be done on the day of LC/MS instrument running for the best quality results.

**CAUTION** Acetonitrile and 2-propanol are toxic. Wear proper personal protective equipment and follow all applicable chemical safety procedures when handling 2- propanol and acetonitrile.

1. 10. Vortex all reconstituted samples at 4 °C for 10 minutes
2. 11. Centrifuge all samples at 30,000g and 4 °C for 15 minutes.
3. 12. Transfer around 40 μL of the supernatant to autosampler glass vials. Choose a consistent volume to be transferred for all samples.

**CRITICAL** For choice of autosampler glass vials, either use glass vials designed for <2 mL volume, or use 2 mL glass vials with preplaced glass inserts. 2 mL glass vials without preplaced glass inserts will not reach an appropriate height for injection into the LC/MS.

#### Preparing pooled quality controls (QCs) for samples

**TIMING** 30 min

**CRITICAL** Pooled quality controls (QCs) are important in lipidomic studies for ensuring reliability of the data being acquired. Separate QCs should be prepared for varying biological environments, such as WC/whole-tissue and LysoIP fractions. QCs should be prepared by acquiring a small amount of the supernatant from each of the replicates.

Prepare undiluted, 1:3, and 1:10 QCs from the pooled mixes.

1. 13. Transfer 5 μL from each autosampler glass vial into a separate pooled QC vial, with one pooled QC vial per biological group. Repeat the same process for any alternative biological groups that need separate pooled mixes. The exact volume aliquoted can be different for groups based on the liquid volume of sample available. **CRITICAL** WC/whole-tissue and LysoIP samples always need separate pooled QCs; however, differences like genotype and treatment conditions within the WC/whole-tissue and LysoIP groups are not necessary to separate in the pooled QCs.
2. 14. Prepare 1:3 and 1:10 QC dilutions for samples by diluting the pooled mixes from the previous step with the appropriate volume of 13:6:1 ACN:IPA:H2O (v/v/v) lipidomic buffer.

**CRITICAL** At the end, there should be QCs for each distinct set of biological samples, all in autosampler glass vials that were pre-loaded with the glass inserts.

**CAUTION** Acetonitrile and 2-propanol are toxic. Wear proper personal protective equipment and follow all applicable chemical safety procedures when handling 2- propanol and acetonitrile.

#### LC/MS with Orbitrap mass spectrometer

**TIMING** 45 min per sample per MS polarity during Orbitrap usage

1. 15. Prepare several glass vials containing only a set volume of the 13:6:1 ACN:IPA:H2O (v/v/v) lipidomic buffer as blank extracts to account for background signals. **CRITICAL** These “blanks” should also be used intermittently during the LC/MS run to wash the needle between samples. Generally, inject a blank sample after every 4 sample injections.

**CAUTION** Acetonitrile and 2-propanol are toxic. Wear proper personal protective equipment and follow all applicable chemical safety procedures when handling 2- propanol and acetonitrile.

1. 16. Connect the lipidomic Mobile Phase A (MPA) bottle to line A and Mobile Phase B (MPB) bottle to line B, respectively, in the LC/MS equipment. See “Reagent Setup” for instructions on preparing these buffers.

**CAUTION** MPA and MPB contain chloroform, 2-propanol, and acetonitrile, which are toxic. Wear proper personal protective equipment and follow all applicable chemical safety procedures when handling MPA and MPB.

1. 17. Load all glass vials to the autosampler (including biological samples, pooled QCs,

*CLN5* KO and WT BMPII standards, and blanks).

1. 18. Set the Vanquish HPLC and Orbitrap instrument parameters as described in the “Equipment Setup” section.
2. 19. Run the instrument with a 4 μL injection volume and monitor real-time LC/MS results.

**PAUSE POINT** After running the instrument, data may be analyzed later as necessitated.

#### Data analysis for Orbitrap mass spectrometer

**TIMING** 1 day

1. 20. Collect the raw data and search the raw data for lipids using LipidSearch (Thermo Fisher Scientific), according to the LipidSearch parameters below. 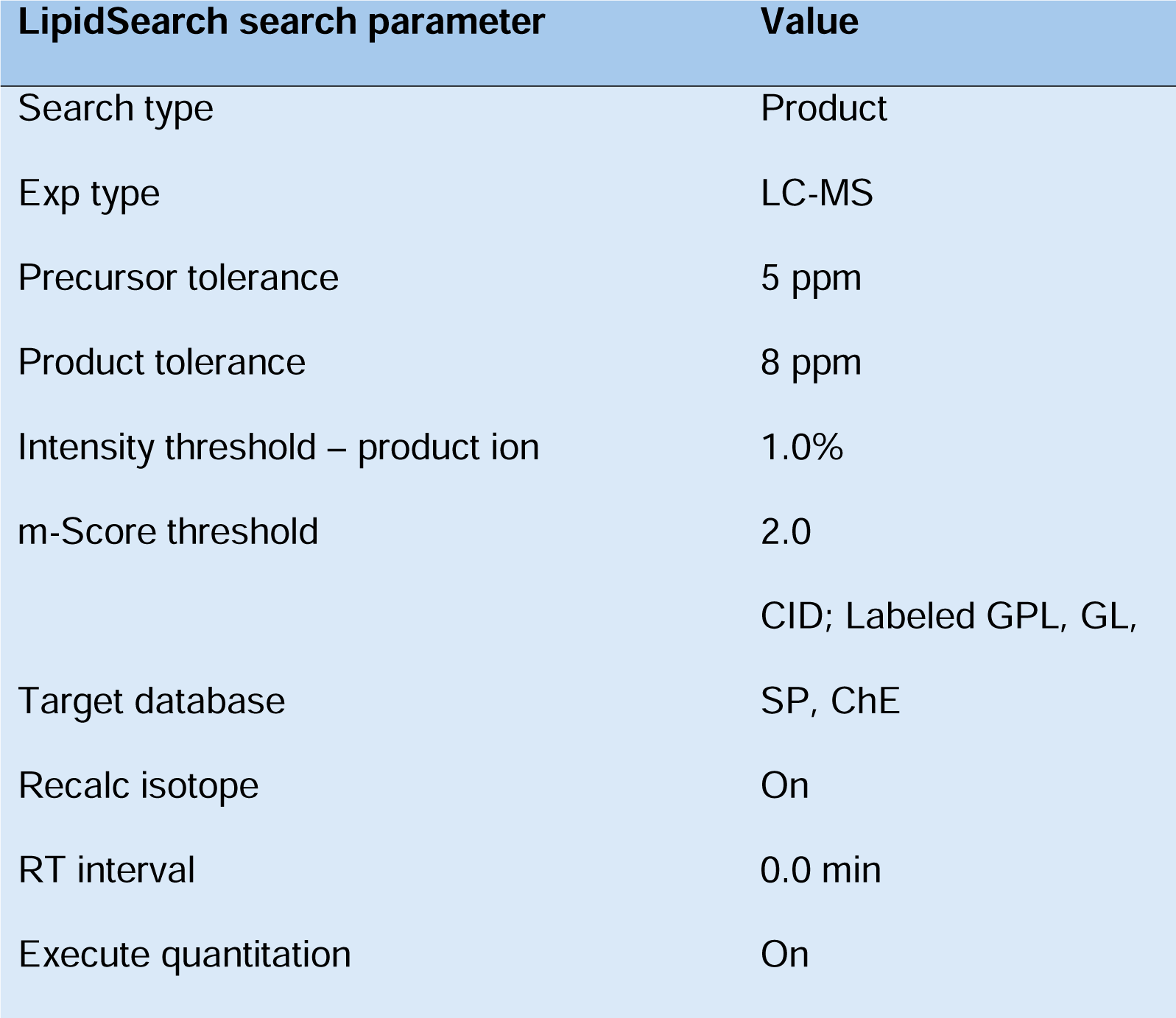
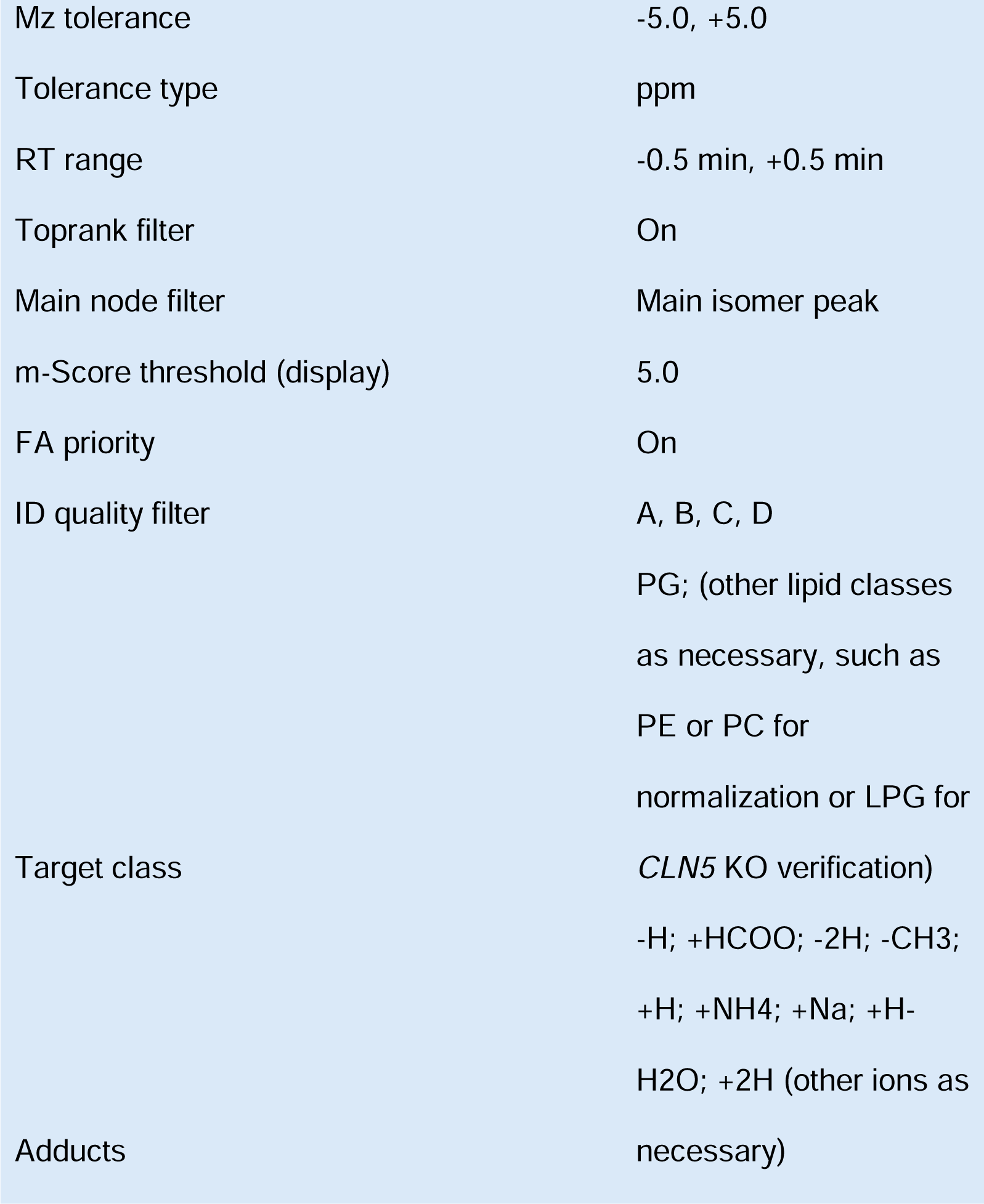
2. 21. Further process the LipidSearch by aligning the search files, according to the LipidSearch parameters below. 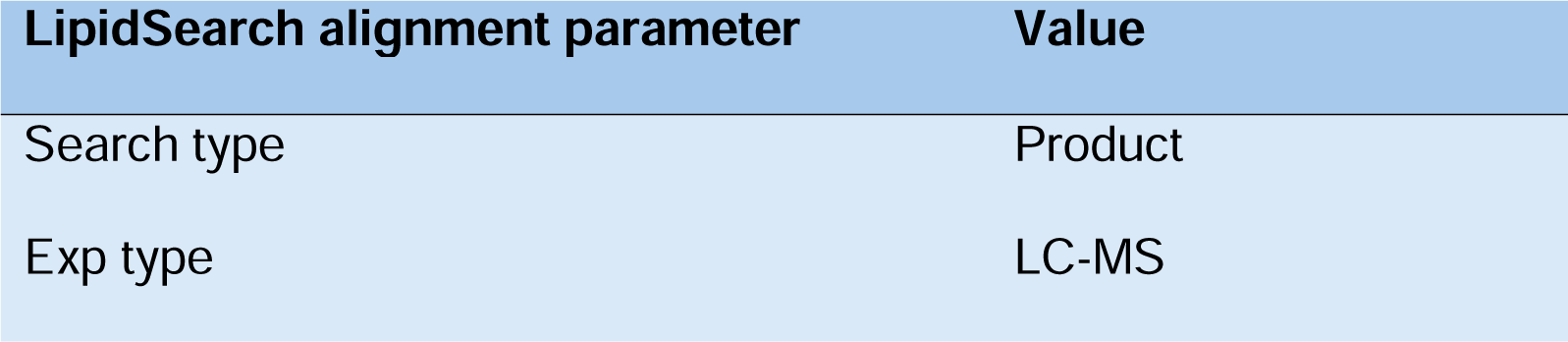
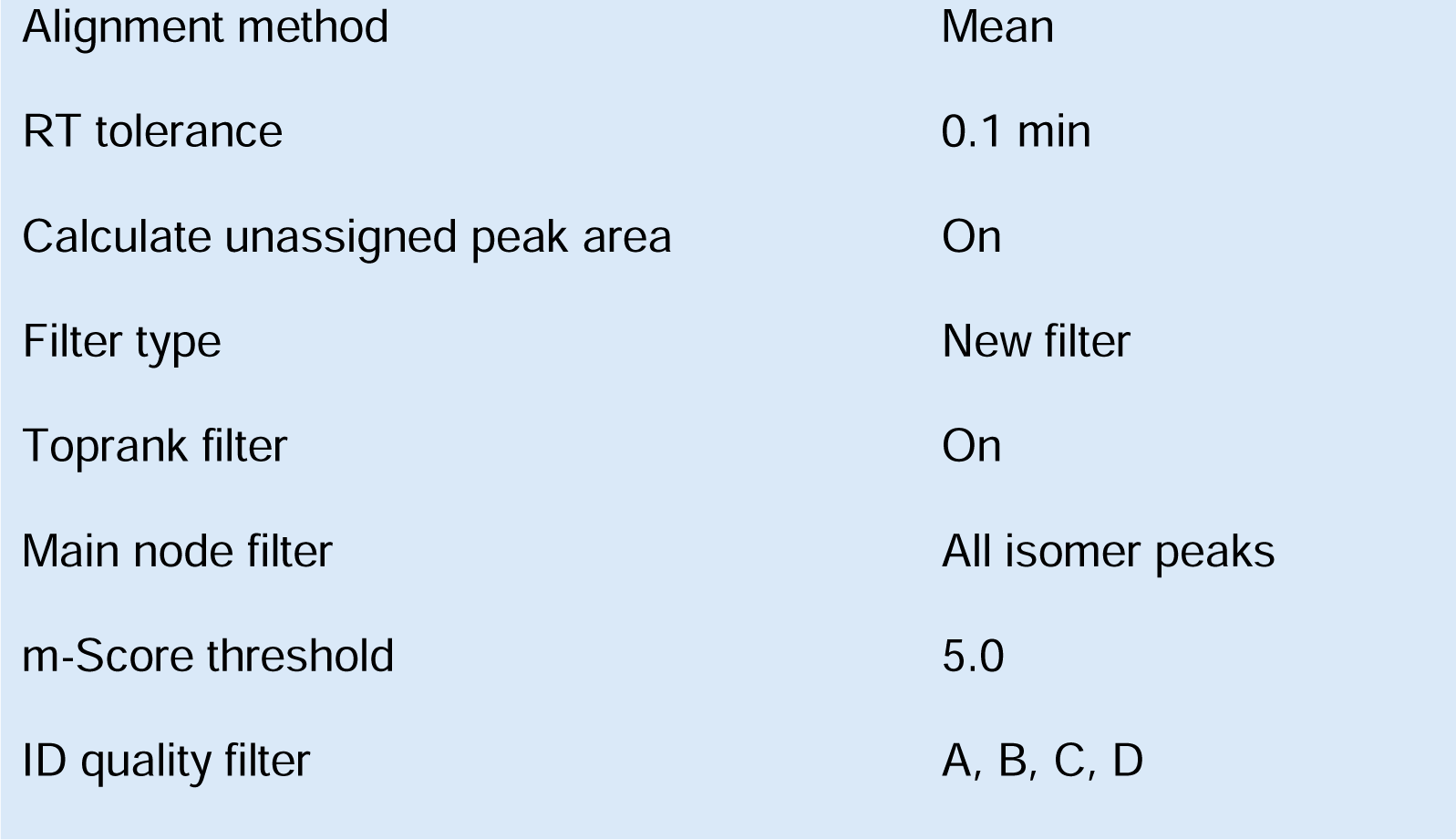
3. 22. Distinguish BMPs and PGs by retention time (RT) differences (Fig. 1A) and by examining the MS2 spectra collected by LipidSearch. The appearance of monoacylglycerol (MG) fragments ([MG-H_2_O+H]^+^) from the positive polarity ammonium adduct ([M+NH_4_]^+^) of PG-annotated species demonstrates that these are truly BMP species. By contrast, the appearance of diacylglycerol (DG) fragments ([DG-H_2_O+H]^+^) from the positive polarity ammonium adduct ([M+NH_4_]^+^) of PG- annotated species demonstrates that these are truly PG species (Fig. 1B, Fig. 2). The characteristic MG and DG fragments for common BMP and PG species are tabulated below, respectively. The characteristic fragments for other species can be derived from their acyl chain compositions and exact isotopic masses. Additionally, to distinguish between acyl chain isomers of PG, check the MS2 spectra of negative polarity [M-H]^-^ ions (Fig. S2). Though these negative ion MS2 spectra will not give the characteristic MG or DG fragments to distinguish between BMP and PG, it will clearly distinguish the two acyl chains of a BMP or PG as fatty acid fragments, eliminating ambiguity as to PG’s acyl chains.

| Species | Parental $m/z$<br>$[M+NH_4]^+$ | Characteristic<br>MG fragment 1<br>$[MG-H_2O+H]^+$ | Characteristic<br>MG fragment 2<br>$[MG-H_2O+H]^+$ |
| --- | --- | --- | --- |
| BMP(14:0_18:1) | 738.5280 | 285.2424 | 339.2894 |
| BMP(16:0_16:0) | 740.5436 | 313.2737 | N/A |
| BMP(16:0_16:1) | 738.5280 | 313.2737 | 311.2581 |
| BMP(16:0_18:0) | 768.5749 | 313.2737 | 341.3080 |
| BMP(16:0_18:1) | 766.5593 | 313.2737 | 339.2894 |
| BMP(16:0_18:2) | 764.5436 | 313.2737 | 337.2737 |
| BMP(16:0_20:4) | 788.5436 | 313.2737 | 361.2737 |
| BMP(16:0_22:6) | 812.5436 | 313.2737 | 385.2737 |
| BMP(16:1_18:1) | 764.5436 | 311.2581 | 339.2894 |
| BMP(16:1_20:3) | 788.5436 | 311.2581 | 363.2894 |
| BMP(16:1_22:6) | 810.528 | 311.2581 | 385.2737 |
| BMP(18:0_18:1) | 794.5906 | 341.3050 | 339.2894 |
| BMP(18:1_18:1) | 792.5749 | 339.2894 | N/A |
| BMP(18:1_18:2) | 790.5593 | 339.2894 | 337.2737 |
| BMP(18:1_20:4) | 814.5593 | 339.2894 | 361.2737 |
| BMP(18:1_22:6) | 838.5593 | 339.2894 | 385.2737 |
| BMP(18:2_18:2) | 788.5436 | 337.2737 | N/A |
| BMP(18:2_20:3) | 814.5593 | 337.2737 | 363.2894 |
| BMP(18:2_20:4) | 812.5436 | 337.2737 | 361.2737 |
| BMP(18:2_22:6) | 836.5436 | 337.2737 | 385.2737 |
| BMP(20:4_20:4) | 836.5436 | 361.2737 | N/A |
| BMP(20:4_22:6) | 860.5436 | 361.2737 | 385.2737 |
| BMP(20:5_22:6) | 858.528 | 359.2581 | 385.2737 |
| BMP(22:4_22:6) | 888.5749 | 389.3050 | 385.2737 |
| BMP(22:5_22:5) | 888.5749 | 387.2894 | N/A |
| BMP(22:5_22:6) | 886.5593 | 387.2894 | 385.2737 |
| BMP(22:6_22:6) | 884.5436 | 385.2737 | N/A |
| BMP(15:0_18:1-d7) | 759.5876 | 299.2581 | 346.3333 |

| Species | Parental $m/z$<br>[M+NH<sub>4</sub>]<sup>+</sup> | Characteristic DG<br>fragment<br>[DG-H<sub>2</sub>O+H]<sup>+</sup> |
| --- | --- | --- |
| PG(14:0_18:1;<br>16:0_16:1) | 738.5280 | 549.4877 |
| PG(16:0_16:0) | 740.5436 | 551.5034 |
| PG(16:0_18:0) | 768.5749 | 579.5347 |
| PG(16:0_18:1) | 766.5593 | 577.5190 |
| PG(16:0_18:2) | 764.5436 | 575.5034 |
| PG(16:0_20:4;<br>16:1_20:3;<br>18:2_18:2) | 788.5436 | 599.5034 |
| PG(16:0_22:6) | 812.5436 | 623.5034 |
| PG(16:1_18:1) | 764.5436 | 575.5034 |
| PG(16:1_22:6) | 810.528 | 621.4878 |
| PG(18:0_18:1) | 794.5906 | 605.5503 |
| PG(18:1_18:1) | 792.5749 | 603.5347 |
| PG(18:1_18:2) | 790.5593 | 601.51904 |
| PG(18:1_20:4;<br>18:2_20:3) | 814.5593 | 625.5190 |
| PG(18:1_22:6) | 838.5593 | 649.51904 |
| PG(18:2_20:4) | 812.5436 | 623.5034 |
| PG(18:2_22:6;<br>20:4_20:4) | 836.5436 | 647.5034 |
| PG(20:4_22:6) | 860.5436 | 671.5034 |
| PG(20:5_22:6) | 858.528 | 669.4877 |
| PG(22:4_22:6;<br>22:5_22:5) | 888.5749 | 669.5347 |
| PG(22:5_22:6) | 886.5593 | 697.5190 |
| PG(22:6_22:6) | 884.5436 | 695.5034 |
| PG(15:0_18:1-d7) | 759.5876 | 570.5473 |

**Figure 2.**
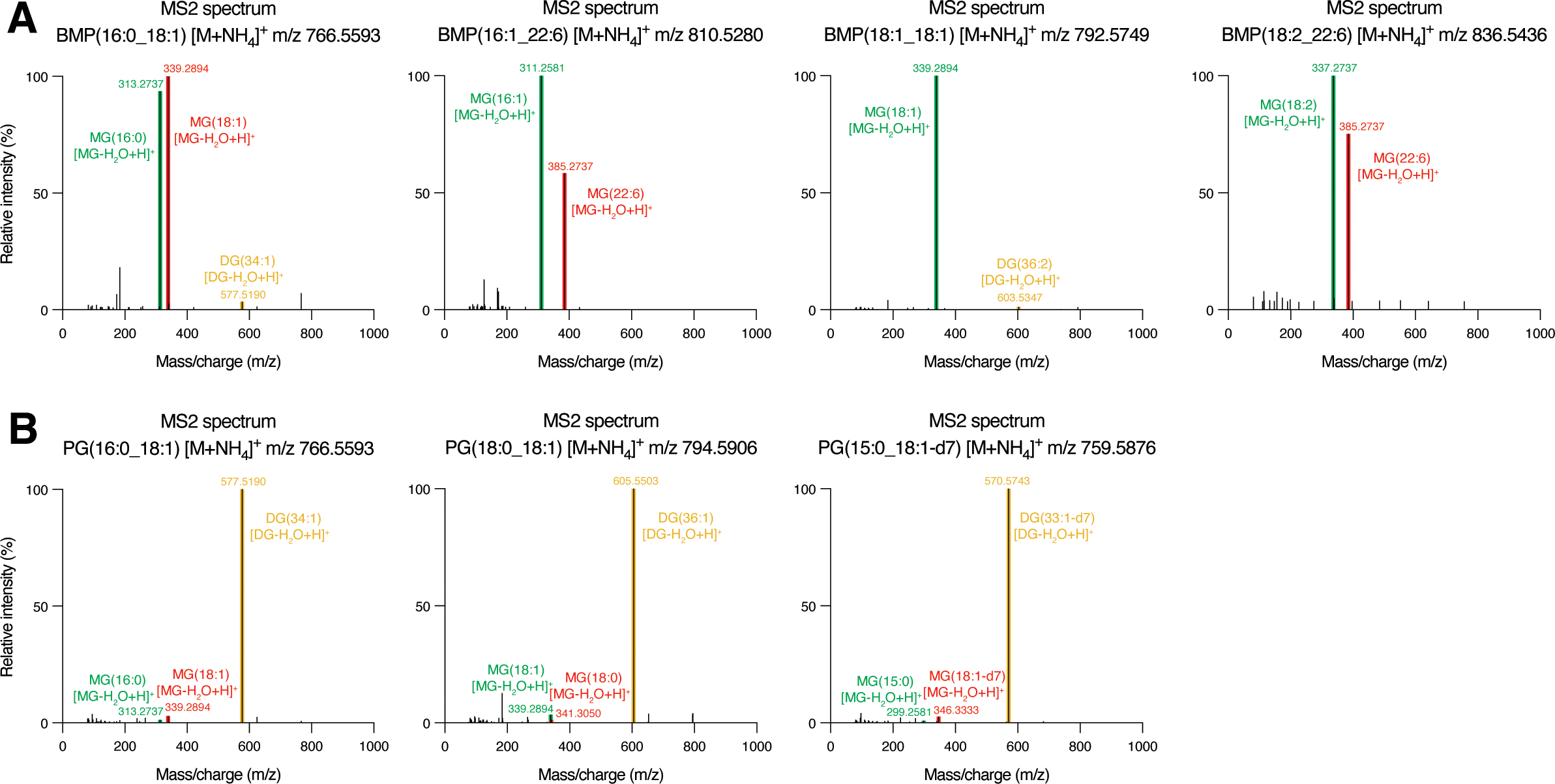
Differential tandem mass (MS2) spectra for BMPs and PGs collected on the Orbitrap LC/MS. A. High-resolution mass spectra for representative BMPs. B. High-resolution mass spectra for representative PGs. MS2 spectra were initially generated and visualized using LipidSearch. MS2 raw data were then extracted from FreeStyle. Lipid fragment annotations are notated from LipidSearch. Relative intensity (%) is relative to highest intensity response. All MS2 presented are from positive mode [M+NH_4_]^+^. All MS2 are from the Orbitrap raw data files and experiment corresponding to Fig. 3B. MG = monoacylglycerol; DG = diacylglycerol.

**CRITICAL** Distinguishing BMPs and PGs based on RT and [M+NH_4_]^+^ MS2 is crucial for confidently annotating BMPs definitively. If RT and MS2 do not provide enough annotation information, then the use of *CLN5* KO and WT BMPII standards is necessary for confident annotation. We suggest using BMPII standards in all cases, for consistent confident annotation.

1. 23. Further distinguish BMPs and PGs using the biological metrics established in this method: BMPII, BMPES, or both (Fig. 1C-D, Fig. 3A-B, Fig. 4A). Specifically, use BMPII metrics from WT and *CLN5* KO BMPII standards, or BMPES metrics from LysoIP and WC samples.

**Figure 3.**
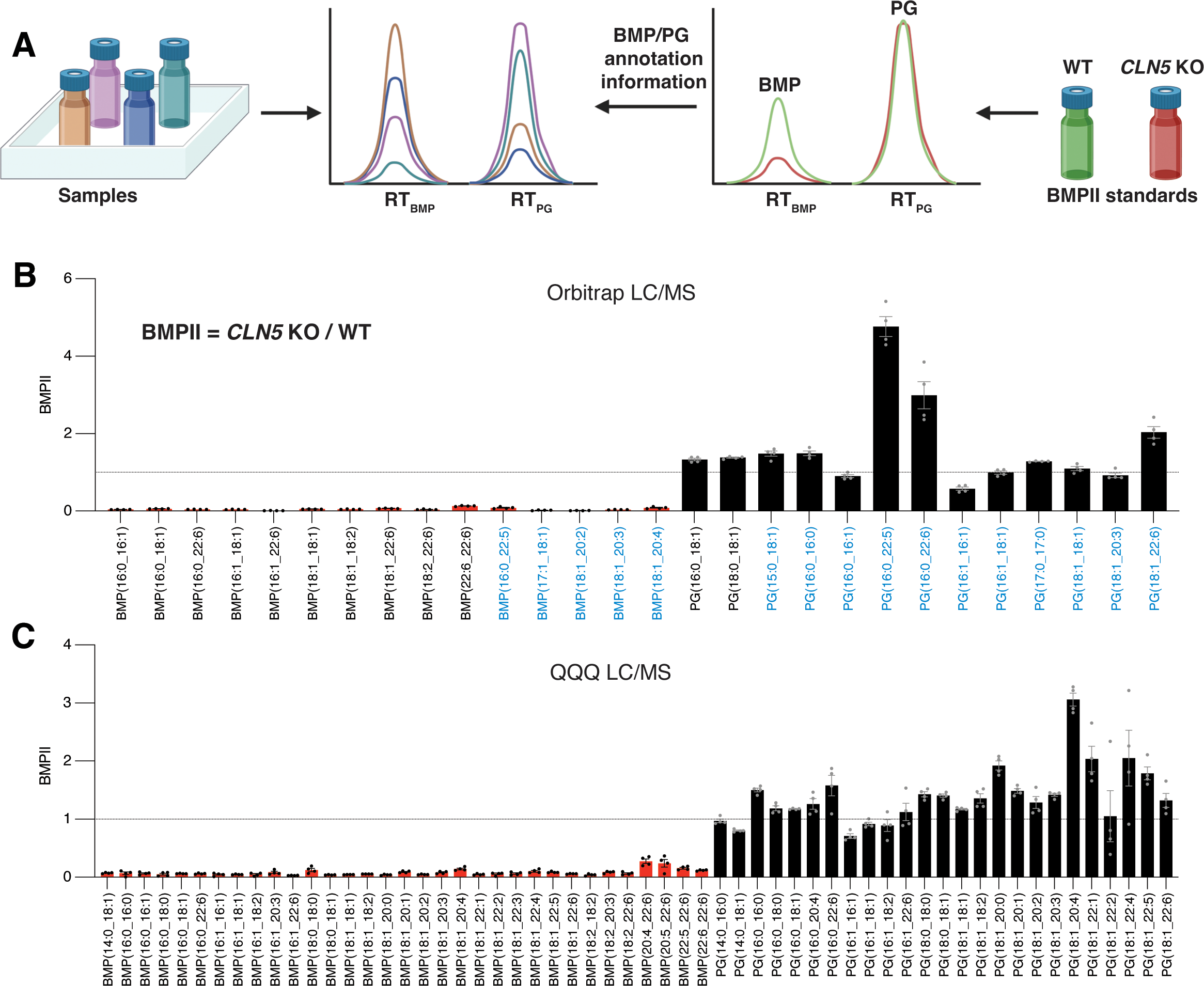
BMP identification index (BMPII) for improved BMP annotations. A. Schematic of applying BMPII for improved BMP annotations. Both BMPII standards and actual samples are analyzed within a single sequence. BMP peaks show a significant reduction in the *CLN5* KO standard, whereas PG peaks remain relatively unchanged. The retention time (RT) information from the BMPII standards is then used to assist in annotating BMPs in the actual samples, distinguishing them from PGs. B. BMPII for BMP (red) and PG (black) species acquired on the Orbitrap LC/MS. Black color BMP and PG names indicate species with definitive +NH_4_ MS2. Blue color BMP and PG names indicate species without definitive +NH_4_ MS2. C. BMPII for BMP (red) and PG (black) species acquired on the QQQ LC/MS. Lipidomics data from whole-cell lipid extracts of wild-type (WT) and *CLN5* knockout (KO) HEK293T cells. Data is presented as mean ± SEM (n = 4). BMPII calculated relative to mean of WT samples. Data from Orbitrap normalized by PE(18:1_18:1). Data from QQQ normalized by PC(16:0_18:1). Dotted line is presented at BMPII = 1.

**Figure 4.**
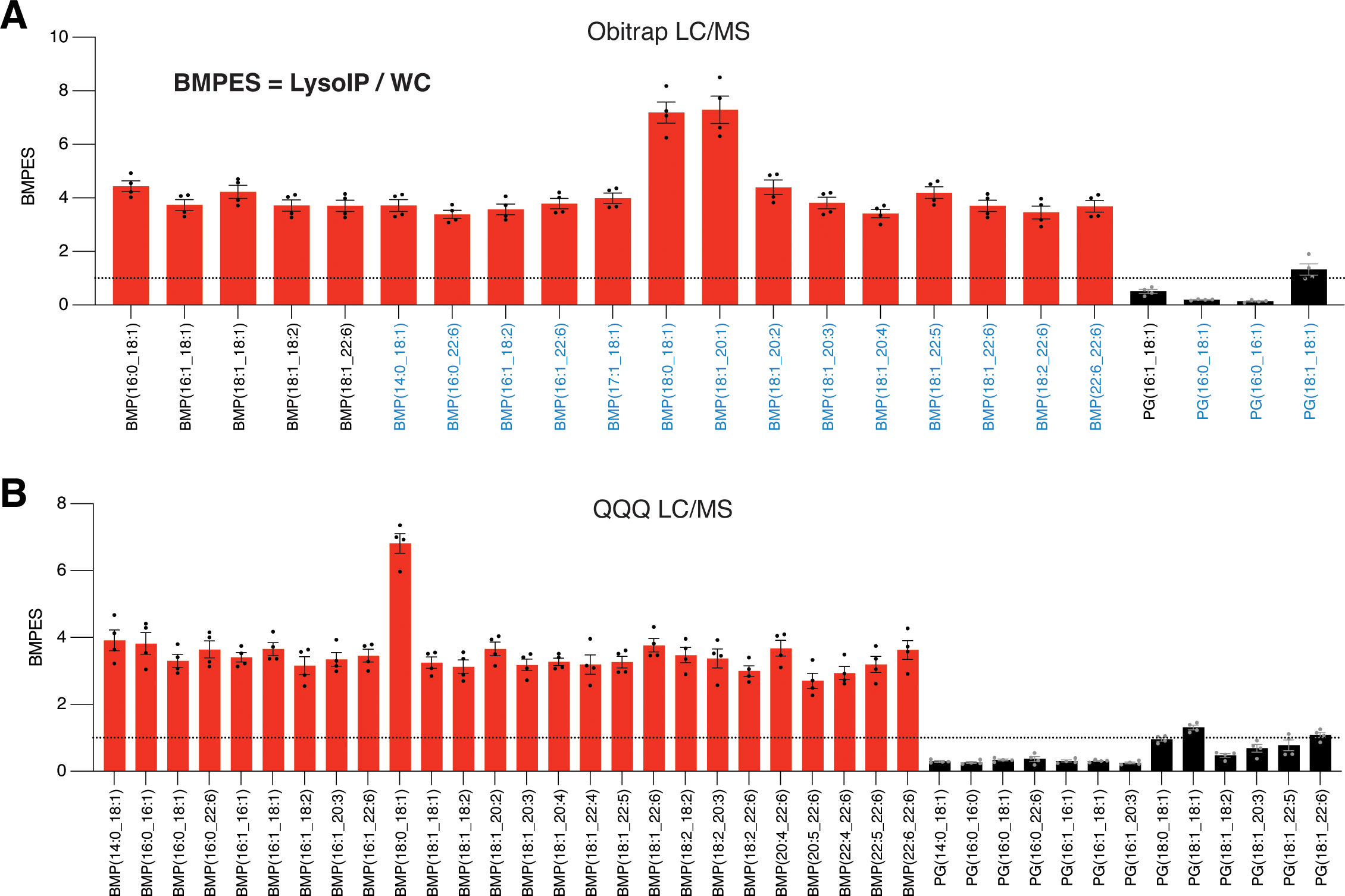
BMP enrichment score (BMPES) for additional improved BMP annotations. A. BMPES for BMP (red) and PG (black) species acquired on the Orbitrap LC/MS. Black color BMP and PG names indicate species with definitive +NH_4_ MS2. Blue color BMP and PG names indicate species without definitive +NH_4_ MS2. B. BMPES for BMP (red) and PG (black) species acquired on the QQQ LC/MS. Lipidomics data from whole-cell and LysoIP lipid extracts of WT SH-SY5Y cells. Data is presented as mean ± SEM (n = 4). BMPES calculated relative to mean of WC samples. Data from Orbitrap normalized by PE(18:1_18:1). Data from QQQ normalized by PC(18:1_18:1). Dotted line is presented at BMPES = 1.

**CRITICAL** Distinguishing BMPs and PGs based on the BMPII and BMPES metrics is crucial for confidently annotating BMPs and PGs that lack definitive [M+NH_4_]^+^ MS2 spectra. BMPII standards are especially valuable, and give strong confidence in the annotation of BMPs if depleted in *CLN5* KO.

1. 24. Optionally, further visualize Orbitrap spectra using FreeStyle to distinguish BMPs and PGs and assist in subsequent TraceFinder method creation.
2. 25. Using TraceFinder, create a method that includes all BMPs, PGs, and other lipids of interest with RTs based on the annotation information from LipidSearch and Compound Discoverer.

**CRITICAL** BMPs and PGs should be quantified using their negative polarity [M-H]^-^ ions. Although ammonium adduct MS2s are necessary for distinguishing BMPs and PGs, the -H negative ion yields the best quantitative results.

1. 26. Using TraceFinder, apply the created method to integrate peak areas.
2. 27. Manually correct peak integrations, ensuring that all integrations accurately include full peaks at the correct RTs.
3. 28. Export raw abundances from TraceFinder for further analysis.
4. 29. Ensure sufficiently high signal-to-noise ratio between samples and blank samples.

Set an established minimum signal-to-noise ratio and apply that threshold to all lipids. If lipids fall below the threshold, exclude them from analysis.

1. 30. Ensure quantitative linearity of target species results using diluted pooled QC samples (Fig. S3).

**CRITICAL** Fold changes of lipid species, including BMPs, PGs, and normalizing lipid species, cannot be accurately reported without ensuring quantitative linearity of the mass spectrometry output. If lipids have poor linear R^2^ using the diluted pooled QC samples, it is indicative that the lipid cannot be reliably quantified and reported.

1. 31. Normalize sample peak areas. Samples can be normalized using isotopically labeled internal standards present in the SPLASH LIPIDOMIX. For BMP and PG, the best choice of internal standard from the SPLASH LIPIDOMIX is PG(15:0_18:1-d7). Samples can also be normalized by orthogonal biological metrics, such as Bradford assay, cell counts, or other equivalent methods. Additionally, samples can be normalized using endogenous lipids, such as phosphatidylcholine (PC) and phosphatidylethanolamine (PE), to account for variations in sample preparation, matrix effects, and differences in input materials and instrumental responses^47^. **CRITICAL** Proper normalization of samples is critical to accurately profile lipidomic alterations, particularly in cases where genotype or condition differences can result in different cell counts.

#### LC/MS with triple quadrupole (QQQ) mass spectrometer

**TIMING** 20 min per sample during QQQ usage

1. 32. Prepare several glass vials containing only a set volume of the 13:6:1 ACN:IPA:H2O (v/v/v) lipidomic buffer as blank extracts to account for background signals. **CRITICAL** These “blanks” should also be used intermittently during the LC/MS run to wash the needle between samples. Generally, inject a blank sample after every 4 sample injections.

**CAUTION** Acetonitrile and 2-propanol are toxic. Wear proper personal protective equipment and follow all applicable chemical safety procedures when handling 2- propanol and acetonitrile.

1. 33. Connect the lipidomic Mobile Phase A (MPA) bottle to line A and Mobile Phase B (MPB) bottle to line B, respectively, in the LC/MS equipment. See “Reagent Setup” for instructions on preparing these buffers.

**CAUTION** MPA and MPB contain chloroform, 2-propanol, and acetonitrile, which are toxic. Wear proper personal protective equipment and follow all applicable chemical safety procedures when handling MPA and MPB.

1. 34. Load all glass vials to the autosampler (including biological samples, pooled QCs,

*CLN5* KO and WT BMPII standards, and blanks).

1. 35. Set the 1290 Infinity II HPLC and QQQ instrument parameters as described in the “Equipment Setup” section.
2. 36. Run the instrument with a 4 μL injection volume and monitor real-time LC/MS results.

**PAUSE POINT** After running the instrument, data may be analyzed later as necessitated.

H. Data analysis for triple quadrupole (QQQ) mass spectrometer

**TIMING** 1 day

1. 37. Extract chromatograms from data in Qualitative Analysis (Agilent Technologies) and QQQ Quantitative Analysis (Quant-My-Way) (Agilent Technologies).
2. 38. Using Qualitative Analysis, check MRM information from selected samples and blank samples to validate and ensure that the MRM information is accurate and does not yield false positives in blank samples.
3. 39. Using Qualitative Analysis, check that the retention time (RT) for lipid transitions across selected sample files are aligned with each other. Additionally, check the total ion chromatogram (TIC) for selected files to make sure there is no RT shift between samples during acquisition. If there is RT shift, note down the shifts for use when annotating BMPs and PGs.
4. 40. Using Qualitative Analysis, distinguish BMP and PG species based on RT. Compare the MRM results for isomeric BMPs and PGs to ensure that the BMP has an earlier RT than its isomeric PG (Fig. 1A, Fig. 5).

**Figure 5.**
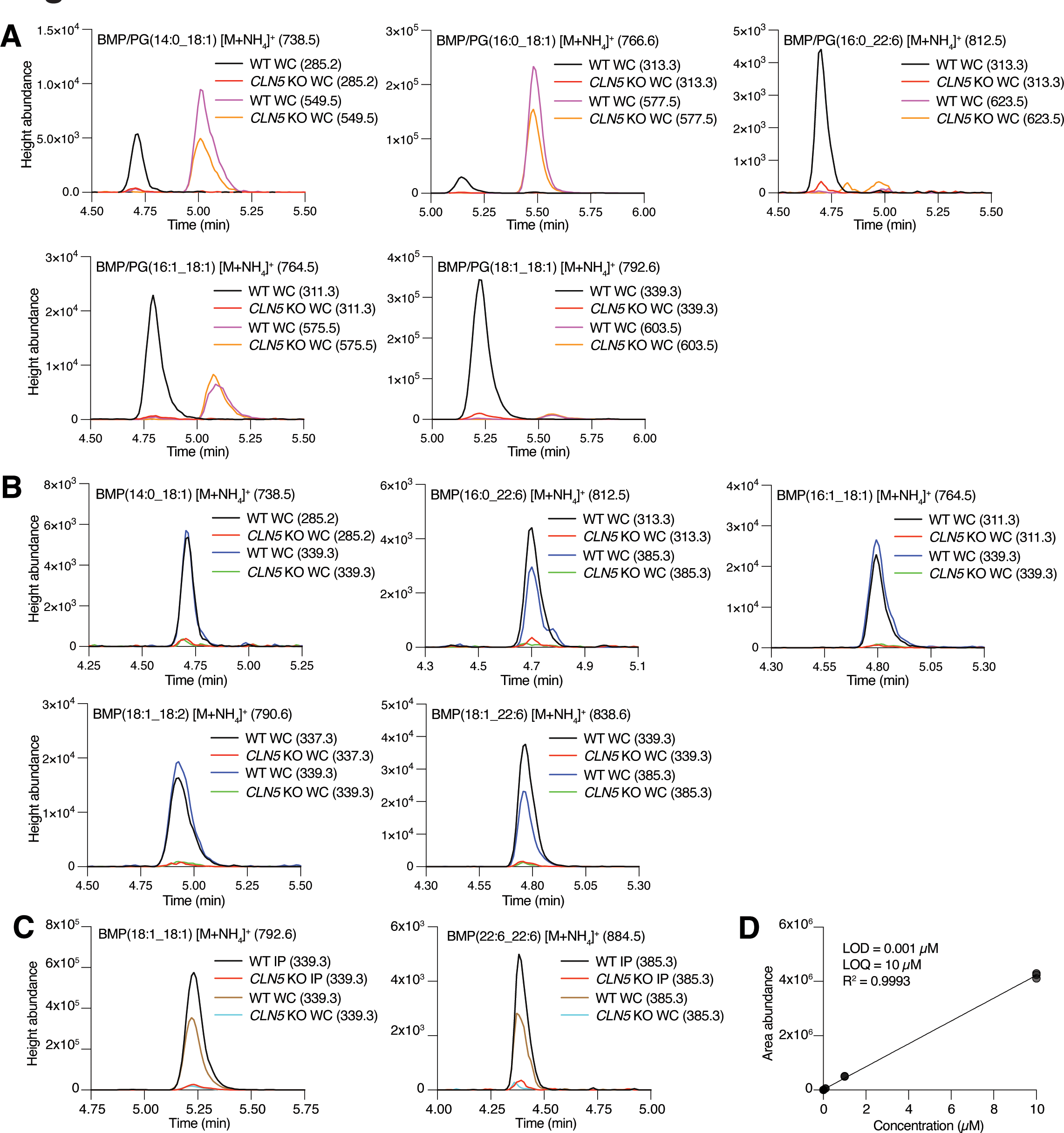
Targeted quantitation for BMPs on the QQQ LC/MS. A. Extracted ion chromatograms for representative BMP (left) and PG (right) species from whole-cell (WC) fractions. B. Extracted ion chromatograms for representative BMPs with two MRM transitions from WC fractions. C. Extracted ion chromatograms for representative BMPs from LysoIP and WC fractions. D. Response linearity check for the BMP(18:1_18:1) standard. Precursor ion mass-to-charge ratio (m/z) in parentheses after lipid name. MRM product ion m/z in parentheses after each respective color and label. All spectra from representative samples within the deposited dataset. Samples are from WC and LysoIP lipid extracts of WT and *CLN5* KO HEK293T cells. Linearity check presented with n = 3. Linearity check R^2^ by simple linear regression. LOD = limit of detection; LOQ = limit of quantitation.

**CRITICAL** Not all acyl chain compositions will have a visible peak for both BMP and PG. In these cases where RTs cannot be compared for isomeric BMPs and PGs, further validation is needed for confident annotation. Due to low abundance fragmentation of BMP into DG and PG into MG, the presence of peaks in either the BMP or PG MRM spectrum cannot be confidently annotated without the corresponding isomeric RT for comparison. In these situations, BMPII metrics from standards of *CLN5* KO and WT, as well as BMPES metrics, can be used to confidently verify BMP or PG annotation.

1. 41. Using Qualitative Analysis, validate that both MRM transitions for BMPs with two transitions yield peaks with matching RTs. If not, these peaks cannot be confidently annotated as BMPs.

**CRITICAL** Ensuring that both BMP transitions yield matching RT peaks is crucial to avoid misannotating non-BMP/PG lipid species, as well as BMP/PG isotopologues, as BMPs.

1. 42. Further distinguish BMPs and PGs using the biological metrics established in this method: BMPII, BMPES, or both (Fig. 1C-D, Fig. 3C, Fig. 4B, Fig. 5A-C). Specifically, use BMPII metrics from WT and *CLN5* KO BMPII standards, or BMPES metrics from LysoIP and WC samples.

**CRITICAL** Distinguishing BMPs and PGs based on the BMPII and BMPES metrics is crucial for confidently annotating BMPs and PGs, especially in cases of low abundance, lack of isomeric RT comparison, or significant non-BMP/PG lipid signals. BMPII standards are especially valuable, and give strong confidence in the annotation of BMPs if depleted in *CLN5* KO.

1. 43. Using Quantitative Analysis, create a method that includes all MRM transitions with RTs corrected based on the annotation information from Qualitative Analysis.
2. 44. Using Quantitative Analysis, apply the created method to integrate peak areas.
3. 45. Manually correct peak integrations, ensuring that all integrations accurately include full peaks at the correct RTs.
4. 46. Export raw abundances from Quantitative Analysis for further analysis.
5. 47. Ensure sufficiently high signal-to-noise ratio between samples and blank samples.

Set an established minimum signal-to-noise ratio and apply that threshold to all lipids. If lipids fall below the threshold, exclude them from analysis.

1. 48. Ensure quantitative linearity of target species results using diluted pooled QC samples (Fig. 5D, Fig. S4).

**CRITICAL** Fold changes of lipid species, including BMPs, PGs, and normalizing lipid species, cannot be accurately reported without ensuring quantitative linearity of the mass spectrometry output. If lipids have poor linear R^2^ using the diluted pooled QC samples, it is indicative that the lipid cannot be reliably quantified and reported.

1. 49. Normalize sample peak areas. Samples can be normalized using isotopically labeled internal standards present in the SPLASH LIPIDOMIX. Samples can also be normalized by orthogonal biological metrics, such as Bradford assay, cell counts, or other equivalent methods. Additionally, samples can be normalized using endogenous lipids, such as phosphatidylcholine (PC) and phosphatidylethanolamine (PE), to account for variations in sample preparation, matrix effects, and differences in input materials and instrumental responses^47^.

**CRITICAL** Proper normalization of samples is critical to accurately profile lipidomic alterations, particularly in cases where genotype or condition differences can result in different cell counts.

### Troubleshooting

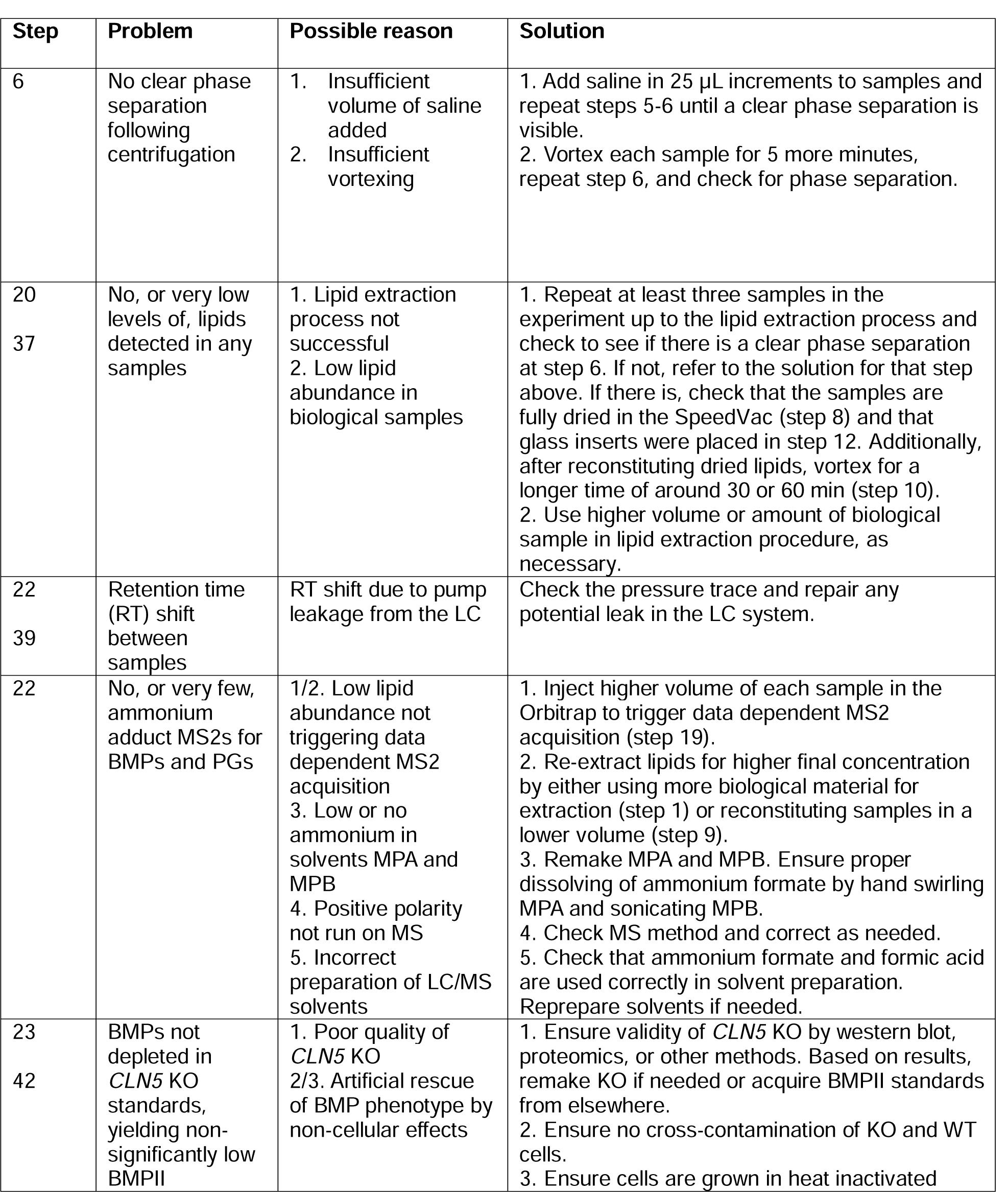
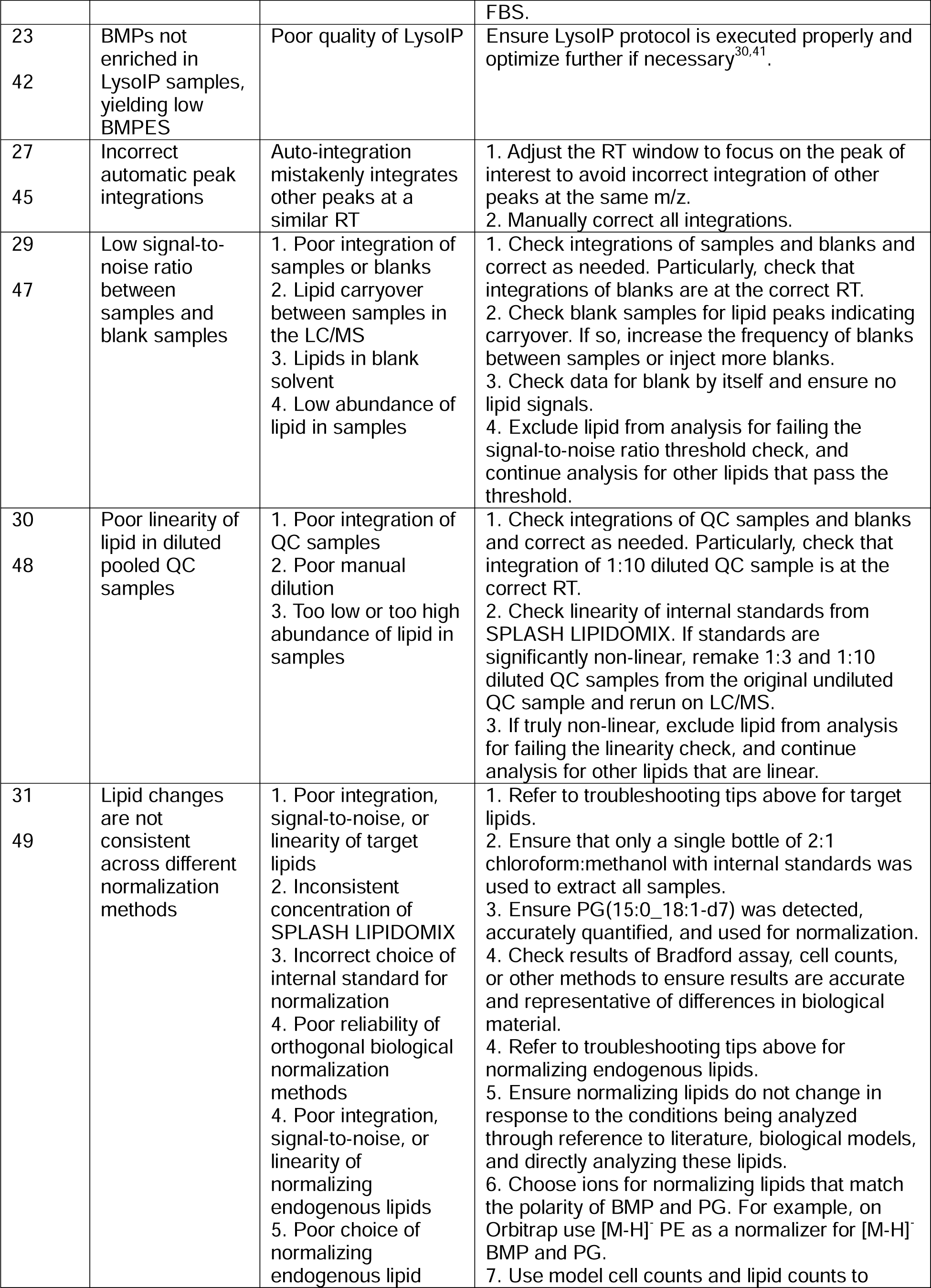
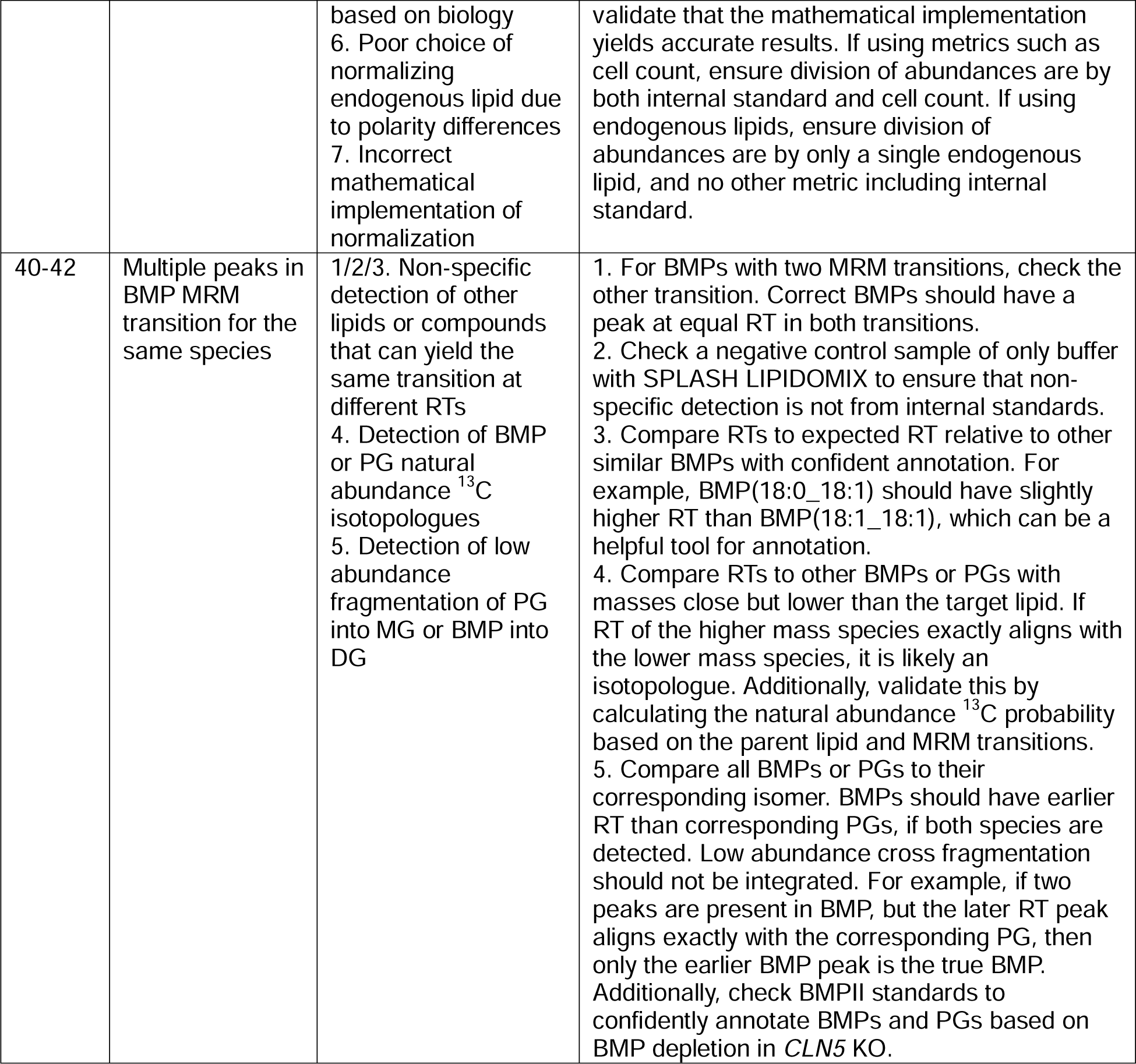

#### Timing

The estimated timing provided below is based on whole-cell harvesting of HEK293T cells from 6-well plates.

Steps 1A(i-x), harvesting of whole-cell samples for BMP analysis, 1 h
Steps 2-12, lipid extraction and processing from harvested samples: 3 h
Steps 13-14, preparing pooled QCs for samples: 30 min
Steps 15-20, LC/MS with Orbitrap: 45 min per sample per MS polarity during Orbitrap usage
Steps 21-27, data analysis for Orbitrap: 1 day
Steps 28-32, LC/MS with QQQ: 20 min per sample during QQQ usage
Steps 33-39, data analysis for QQQ: 1 day

#### Anticipated results

##### Curation of MS2 spectra for representative BMPs and PGs from the Orbitrap LC/MS

Tandem mass spectrometry analysis using positive ionization mode in the presence of ammonium ions results in distinct fragmentation patterns between BMPs and PGs. This allows for clear differentiation between these two isomeric lipid classes. The representative high-resolution mass spectra obtained from the Orbitrap clearly illustrate these differences (Fig. 2).

During positive ionization, fragmentation predominantly occurs at the phosphodiester bonds of the lipids. This process yields distinct fragments characteristic for each lipid class: monoacylglycerol (MG) fragments from BMPs, and diacylglycerol (DG) fragments from PGs. The MG fragments arise from cleavage occurring between phosphate and glycerol on both sides of the lipid, leaving glycerols each with a single acyl chain attached. This yields the signature fragmentation pattern of MGs for BMPs (Fig. 2A). In contrast, the same phosphodiester cleavage of PG results in a glycerol with two acyl chains attached. This yields the signature fragmentation pattern of DG for PGs (Fig. 2B). These specific fragmentation patterns are crucial for accurate lipid classification and can be further substantiated by detailed examination of the MS2 spectra. By referencing a comprehensive table of characteristic fragments, as provided above, one can confidently annotate the tandem mass spectra for BMPs and PGs.

Although MS2 fragmentation from [M+NH_4_]^+^ ions yields the distinct MG or DG fragmentation, MS2 analysis of [M-H]^-^ ions can also provide other valuable information, including acyl chain compositions in the form of fatty acid fragments (Fig. S2). This information is particularly useful for annotation of PGs with only a DG fragment, where acyl chain-dependent isomers, such as PG(18:2_22:6) and PG(20:4_20:4), may not be distinguishable without analysis of negative ionization MS2s. Overall, MS2 analysis gives powerful and unambiguous identification and differentiation of these isomeric lipid classes.

##### Targeted quantitation of BMPs and PGs from the Triple Quadrupole LC/MS

Ammonium adducts of BMPs and PGs were quantified from the QQQ using their characteristic multiple reaction monitoring (MRM) transitions of monoacylglycerol (MG) fragments and diacylglycerol (DG) fragments, respectively. The representative extracted ion chromatograms show that the more polar BMP elutes before its corresponding PG counterpart under the chromatographic conditions (Fig. 5A). As expected, the RTs of BMPs with two MRM transitions, based on their two different acyl chains yielding two distinct MGs, aligned with each other (Fig. 5B). The RTs of BMPs from lysosomal and whole-cell fractions also aligned with each other, despite varying biological matrices (Fig. 5C). Importantly, whole-cell fractions from *CLN5* knockout (KO) cells show significantly depleted BMP levels compared to wild-type (WT) conditions, whereas depletion was not observed for PGs (Fig. 5A-C). To ensure reliable relative quantitation, a quality control linearity check was done using a BMP(18:1_18:1) standard, demonstrating a strong linear response as well as a four order of magnitude difference between the limit of detection (LOD) and limit of quantitation (LOQ) (Fig. 5D). Similar quality control linearity checks were performed for all quantified BMPs and PGs (Fig. S3, Fig. S4).

##### Establishing the BMP identification index (BMPII) and BMP enrichment score (BMPES) for BMP annotations

To further improve BMP annotations, we developed metrics based on two biological characteristics of BMP: 1) BMPs are synthesized by the protein CLN5, and 2) BMPs are predominantly enriched in the endolysosomal pathway. The first metric, the BMP Identification Index (BMPII), was developed to measure the fold change in intensity of candidate BMP peaks in *CLN5* KO cells compared to WT cells. In particular, our method emphasizes the use of BMPII standards, which are standard samples of lipid extracts from WT cells and *CLN5* KO cells that are premade and then run in a single sequence along with experimental samples (Fig. 3A). Due to the biological role of CLN5 in BMP synthesis, a significant reduction in BMP levels in the *CLN5* KO BMPII standard relative to the WT BMPII standards supports the identification of matching peaks in experimental samples as BMPs. Our results clearly demonstrate that the BMPII is a robust identifier for BMPs, showing significant differences in the BMPII between PGs and BMPs on two separate spectrometers, affirming the efficacy of the BMPII metric from BMPII standards (Fig. 3B-C, Fig. 5A, Fig. S1).

The second metric, termed the BMP Enrichment Score (BMPES), measures the relative abundance of candidate BMP peaks in lysosomal compared to whole-cell fractions via LysoIP. The hypothesis is that BMPs, due to their enrichment in the endolysosomal pathway, should exhibit a higher BMPES compared to PGs. Through relative quantitative analysis, our data show that BMPES effectively differentiates BMPs from PGs, albeit with a less pronounced difference than anticipated (Fig. 4). However, BMPES still serves as valid secondary evidence for confirming BMP annotations. Together with the differential RT profiles and MS2 fragmentation patterns, BMPII and BMPES provide additional valuable information for distinguishing BMPs from PGs in complex biological samples. This approach is particularly advantageous in scenarios where the species abundance is too low to trigger data dependent MS2 acquisition. Our results establish that the BMPII and BMPES are reliable metrics that can be seen using both the Orbitrap and QQQ spectrometers, and have been actively in use in our work across a variety of biological systems. By defining the BMPII and BMPES, we establish a robust framework that enhances the reliability of BMP annotations. Our method illustrates the value in combining quantitative biological metrics with detailed spectral analysis for lipidomic profiling.

##### Validating BMP changes in urine samples from patients with the LRRK2 G2019S mutation

As discussed previously, multiple studies have found elevated levels of BMP in urine samples from patients with Parkinson’s disease^23–25^. Gomes et al. (2023) specifically examined samples from patients with mutations and risks in Parkinson’s-associated genes and found that urine BMPs are elevated in patients with the leucine-rich repeat kinase 2 (*LRRK2*) G2019S mutation^25^ (Fig. 6A). To validate these results using our BMP method, we acquired a subset of the control and *LRRK2* G2019S urine samples and used our BMP method with the Ultivo QQQ LC/MS. Our method replicated and validated the BMP elevation in LRRK2, revealing statistically significant elevation of the top three BMPs by abundance: BMP(22:6_22:6), BMP(18:1_22:6), and BMP(18:1_18:1) (Fig. 6B). Furthermore, we quantified all detectable BMP species and showed significant elevation in most BMP species (Fig. 6C). Demonstrating the power of our method, we were able to annotate over 30 BMPs, vastly expanding the number of unique species compared to the previous work. Moreover, our BMP method proved to be robust, effectively analyzing not only common *in vitro* and *in vivo* cell materials but also clinical samples from patients.

**Figure 6.**
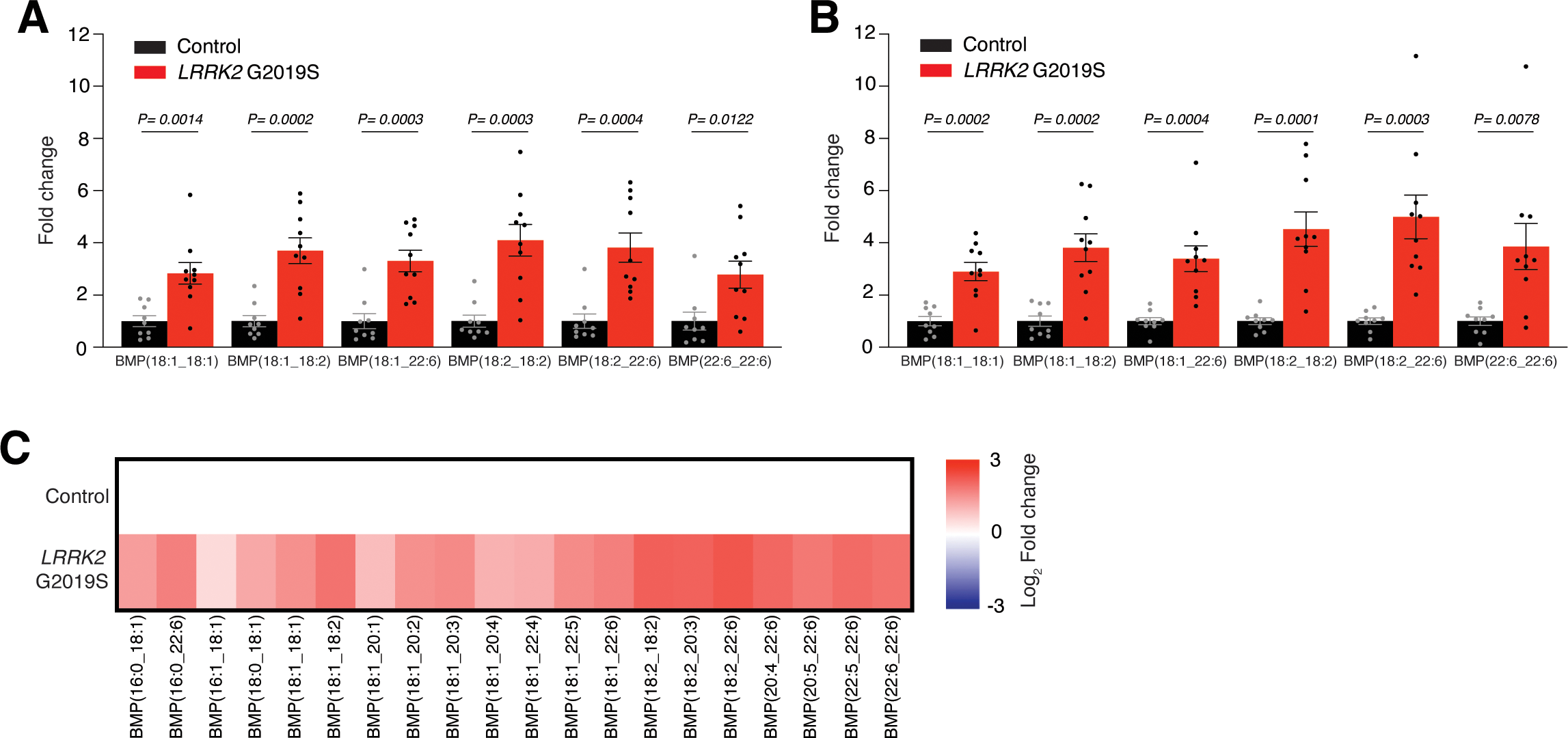
Altered BMP levels in *LRRK2* G2019S patient urine samples. A. Results from Gomes et al., 2023, showing increased BMP in urine of patients with *LRRK2* G2019S mutations, quantified by Nextcea, Inc. B. Targeted quantitation by QQQ LC/MS using our method, reproducing the previously reported results. C. Heatmap showing the change of all detected BMP species in urine of patients with *LRRK2* G2019S, quantified by QQQ LC/MS. Lipidomics data from urine of control patients (n = 9) and patients with *LRRK2* G2019S (n = 10). Data is presented as mean ± SEM. P-values by Student’s t-tests. Lipidomics data presented as fold change and log_2_ fold change relative to mean of control samples. Data from Nextcea, Inc. normalized by creatinine. Data from QQQ normalized by PC(16:0_18:1).

##### Comparison of BMP quantitation results by LC/MS and antibody staining

We quantified BMPs by anti-BMP antibody staining as detailed in previous studies^35,36,48,49^. Utilizing these antibodies, we compared the total BMP levels in whole-cell HEK293T lipid extracts from *CLN5* KO, WT, and recombinant CLN5 protein (rCLN5) rescue cell models. Contrary to expectations, BMP levels were found to be higher in the *CLN5* KO compared to both the WT and rCLN5 rescue (Fig. 7A). This observation is opposite to the anticipated results, considering the known function of CLN5 as a BMP synthase^40^. The loss of CLN5 should result in decreased BMP synthesis, and accordingly, lower BMP levels in the *CLN5* KO samples, which is correctly revealed by our LC/MS-based analysis (Figure 7B). These LC/MS-based results are thus in contrast to those obtained with the antibody-based method. Anti-BMP staining also revealed no change in BMP levels between WT and either *GRN* (*CLN11*) KO or *CLN3* KO, failing to replicate previously reported results of BMP depletion in both *GRN* KO^28,29^ and *CLN3* KO^30–32^ (Fig. S5A). Furthermore, dot blot analysis of lipid standards with the anti-BMP antibody indicated positive signals not only for BMP but for other non-BMP phospholipids as well (Fig. S5B). Notably, the antibody showed staining for lysophosphatidylglycerol (LPG), which is known to be significantly accumulated in *CLN5* KO^40^, possibly explaining the increased stain intensity seen in the *CLN5* KO as compared to WT and rCLN5 rescue. Thus, these results argue against the ability of this commonly used antibody to reliably detect a reduction in BMP levels across various genetic models.

**Figure 7.**
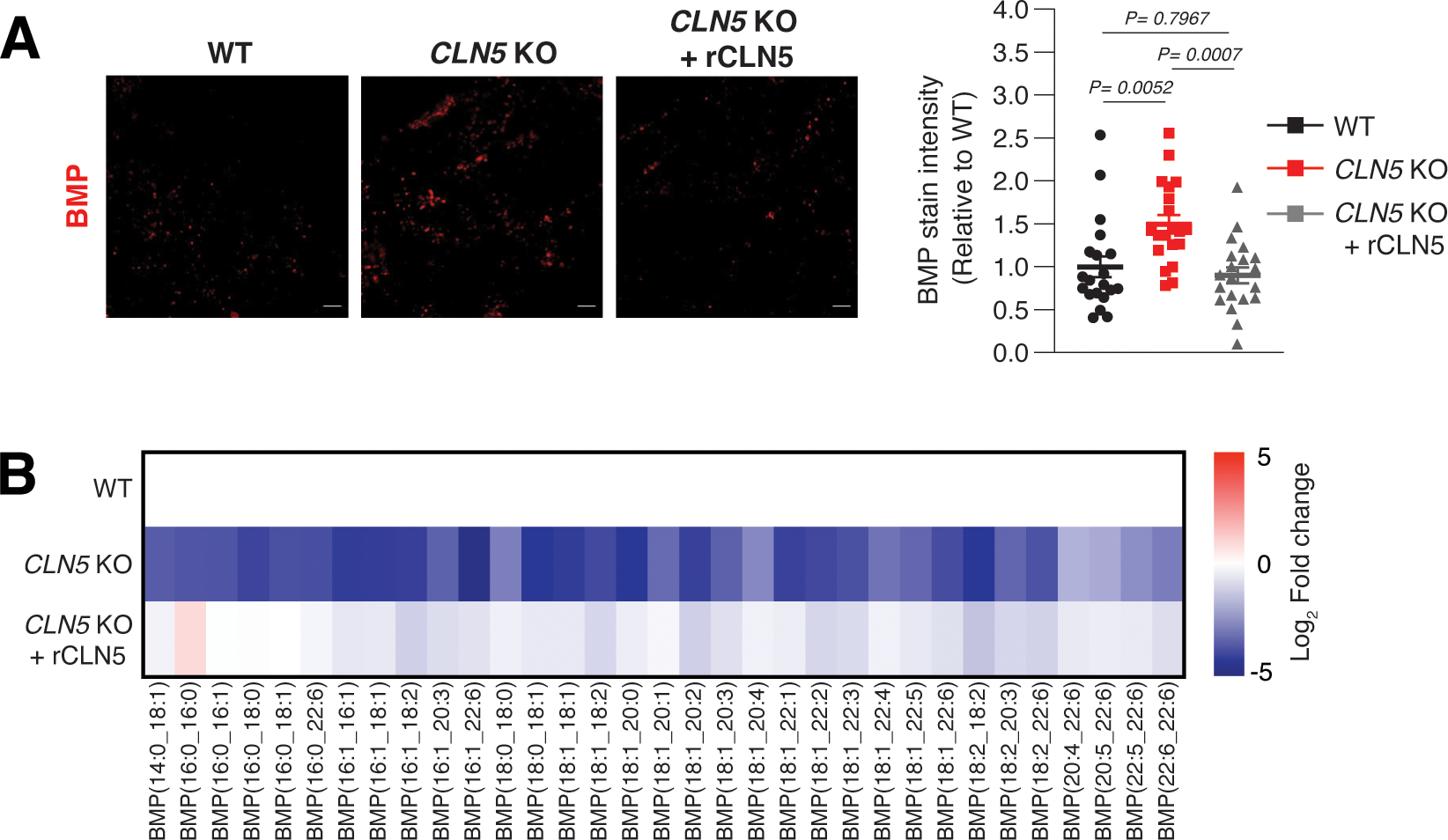
Comparison of BMP profiling by LC/MS and anti-BMP staining. A. BMP levels in *CLN5* KO and rCLN5 rescue cells quantified by anti-BMP antibody. B. BMP levels in *CLN5* KO and rCLN5 rescue cells quantified using our method by QQQ LC/MS. Anti-BMP staining in WT, *CLN5* KO, and *CLN5* KO + recombinant CLN5 protein (rCLN5)-treated HEK293T cells. Images presented with scale bar = 5 µM. Stain intensity data is presented as mean ± SEM (n = 20). P-values by one-way ANOVA with post-hoc Tukey’s HSD test. Lipidomics data presented as log_2_ fold change relative to mean of WT samples. Data from QQQ normalized by PC(16:0_18:1). Lipidomics data from same experiment as Fig. 3B-C, with same data files as Fig. 3C. As in Fig. 4C, lipidomics data from whole-cell lipid extracts of WT, *CLN5* KO, and *CLN5* KO + rCLN5 HEK293T cells.

Collectively, these results demonstrate the reliability of our LC/MS-based method for identifying and quantifying BMPs in biological research, as evidenced by the consistency with the known biochemical functions of CLN5. This contrasts with the deviations observed in BMP antibody staining, further supporting LC/MS as a more dependable method for BMP quantitation than antibody staining. In addition, our LC/MS analysis provides chain-specific information for different BMP species, a level of detail absent in the staining method. This capability enhances our understanding of BMP species, distinguishing LC/MS as a superior analytical tool.

## Concluding remarks

Our LC/MS method allows for robust profiling of BMPs in a variety of biological samples. Using both an Orbitrap LC/MS system and triple quadrupole LC/MS system, we can reliably identify and quantify BMPs to characterize BMP biology and its changes in diseases. We have leveraged conventional LC/MS means to distinguish BMPs and PGs, utilizing differential molecular polarity and MS2 fragmentation. Furthermore, we establish the BMPII and BMPES as metrics for BMP annotation based on the recent biochemical insight of CLN5 synthesis of BMP and the known lysosomal enrichment of BMP, respectively. Unlike other methods for BMP profiling, our approach does not rely on chemical derivatization and still allows for accurate identification and quantitation of specific BMP species. With the established workflow and instrumentation, our method enables fast profiling of BMPs, which we have been able to use to study lysosomal, cellular, whole-cell/tissue, and patient samples. Our methods have paved a confident way forward for precise and comprehensive BMP characterization in health and disease.

## Author contributions

W.D., K.N., E.S.R., and M.A-R. conceived and designed the protocol. W.D., K.N., E.S.R., U.N.M., and M.A-R. conceived and designed experiments. W.D., K.N., E.S.R., U.N.M., J.X., C.C.B., H.N.A., H.Y.L., S.G., T.H., M. A., F.H., and E.S. performed experiments. W.D., K.N., E.S.R., U.N.M., and F.H. performed data analysis. W.D., K.N., E.S.R., C.C.B., and M.A-R. wrote the original draft of the manuscript. All authors read and edited the final manuscript.

## Acknowledgment

We thank all members of the Abu-Remaileh laboratory for helpful insights. We also thank Suzanne Pfeffer, Dario Alessi, and Miratul Muqit for discussion and advice. We would like to thank the Metabolomics Knowledge Center at Stanford Sarafan ChEM-H (directed by Dr. Yuqin Dai). This work was supported by grants from Aligning Science Across Parkinson’s [ASAP-000463] through the Michael J. Fox Foundation for Parkinson’s Research (MJFF) and the Knight Initiative for Brain Resilience to M.A-R. K.N. is supported by the Sarafan ChEM-H Chemistry/Biology Interface Program as a Kolluri Fellow. K.N is additionally supported by the Bio-X Stanford Interdisciplinary Graduate Fellowship affiliated with the Wu Tsai Neurosciences Institute (Bio-X SIGF: Mark and Mary Steven’s Interdisciplinary Graduate Fellow). H.N.A. is supported by the Arc Institute Graduate Fellowship Program. M.A-R is a Stanford Terman Fellow and a Pew-Stewart Scholar for Cancer Research, supported by the Pew Charitable Trusts and the Alexander and Margaret Stewart Trust.

## Data and materials availability

All LC/MS raw data and other raw data will be archived in general purpose repositories with free access following peer review and upon publication.

---

**Figure S1.**
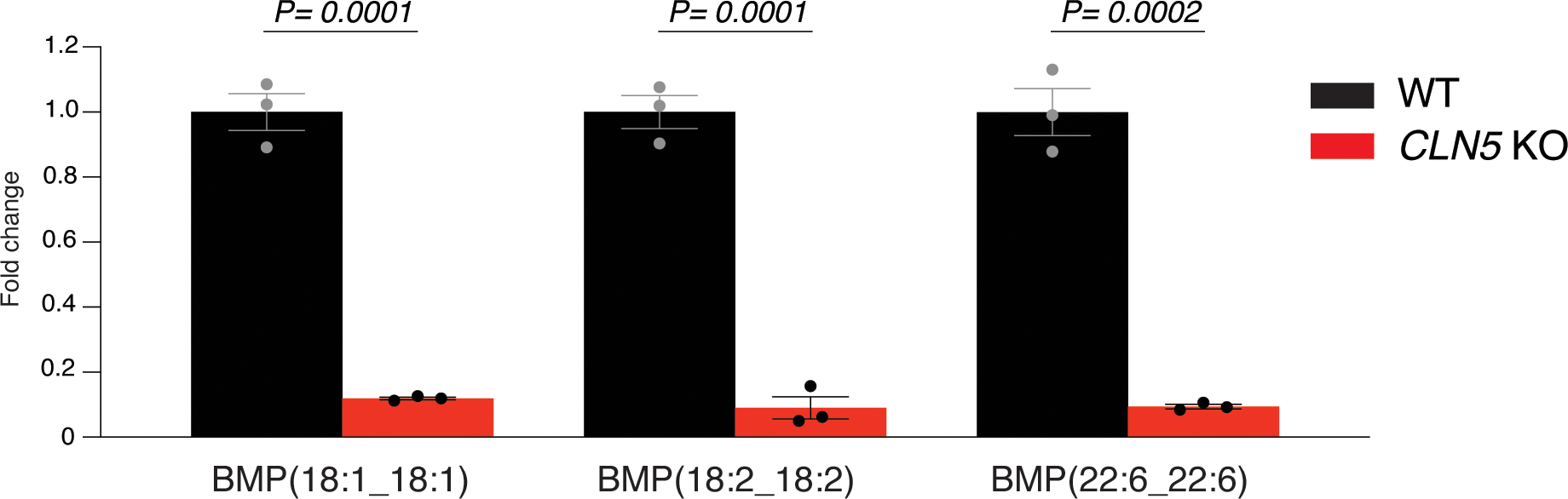
Depleted BMP in *CLN5* KO validated by Nextcea, Inc. Targeted quantitation of BMPs from Nextcea, Inc. show marked reduction of BMP in *CLN5* KO relative to WT. Lipidomics data from whole-cell lipid extracts of WT and *CLN5* KO HEK293T cells. Data is presented as mean ± SEM (n = 3). Fold change calculated relative to mean of WT samples. Data from Nextcea, Inc. normalized by total protein content. BMPs quantified by UPLC-MS/MS using a SCIEX Triple Quad™ 7500 LC-MS/MS System (SCIEX, Framingham, MA) and Shimadzu Nexera XR ultra high-performance liquid chromatograph (UPLC) system (Shimadzu Scientific Instruments, Japan). Corresponding mass-shifted, stable isotope-labeled BMP, di-18:1 BMP-d5 and di-22:6 BMP-d5, employed as internal standards.

**Figure S2.**
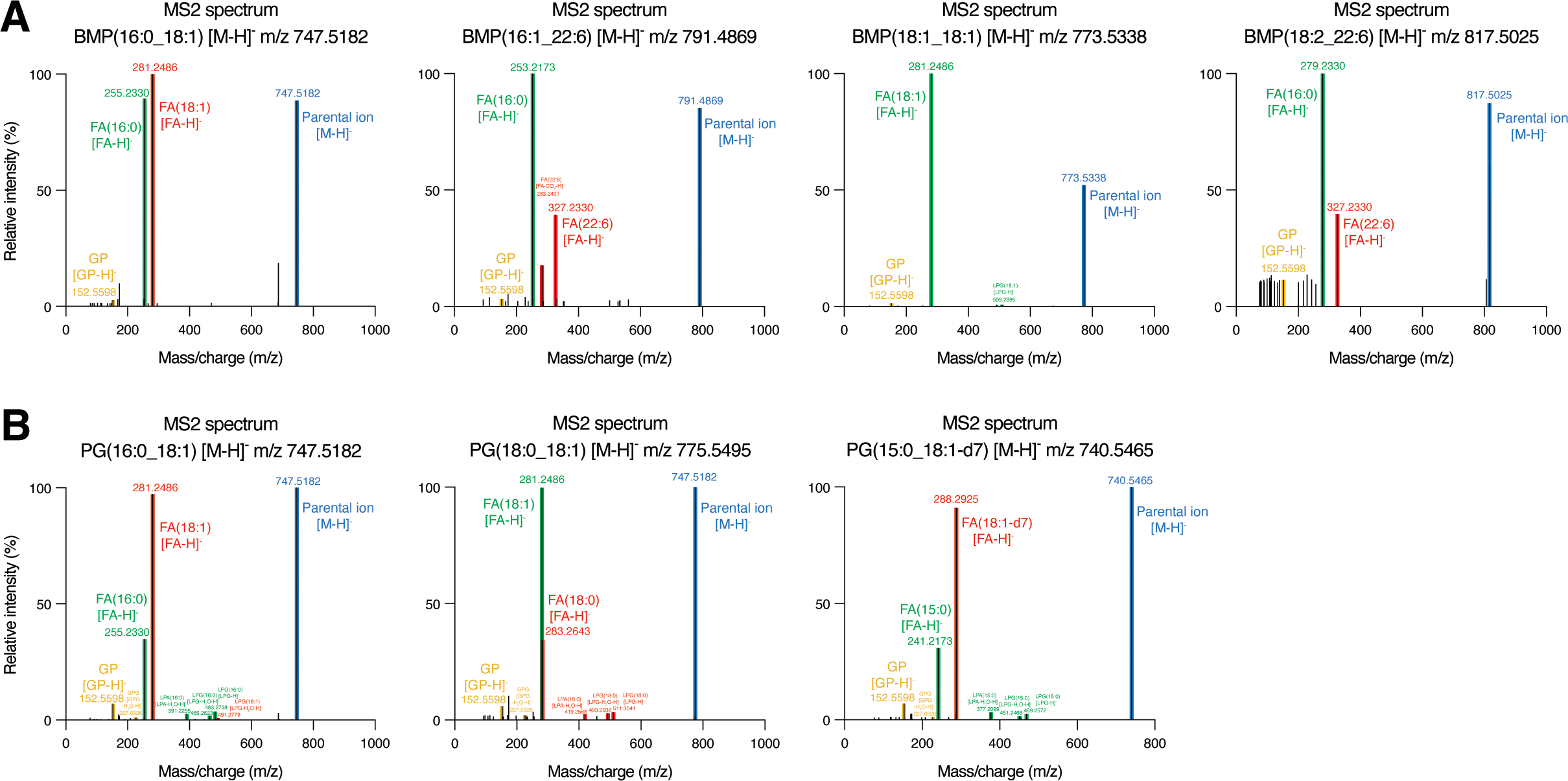
Non-differential negative mode tandem mass (MS2) spectra for BMPs and PGs. A. High-resolution mass spectra for representative BMPs. B. High-resolution mass spectra for representative PGs. MS2 spectra were initially generated and visualized using LipidSearch. MS2 raw data were then extracted from FreeStyle. Lipid fragment are annotated from LipidSearch. Relative intensity (%) is relative to highest intensity response. All MS2 presented are from negative mode [M-H]^-^. All MS2 are from the Orbitrap raw data files and experiment corresponding to Fig. 3B. GP = glycerophosphate; GPG = glycerophosphoglycerol; FA = fatty acid; LPA = lysophosphatidic acid; LPG = lysophosphatidylglycerol.

**Figure S3.**
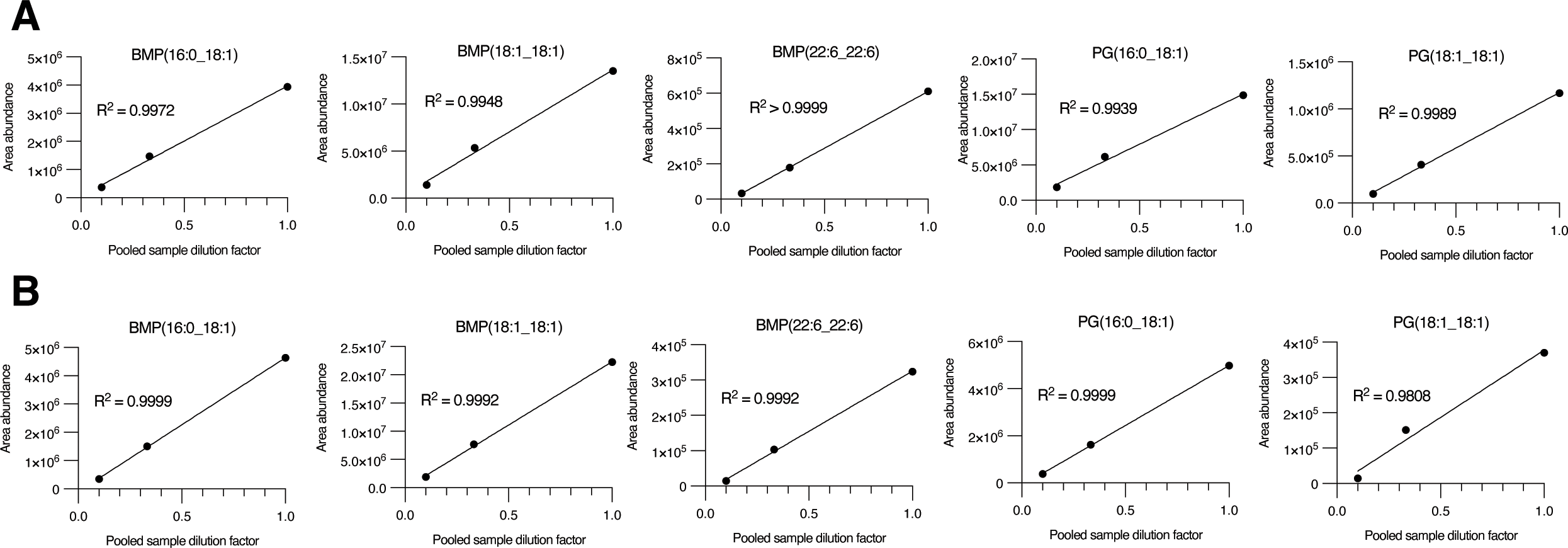
Linearity quality control check for BMP and PG profiling by Orbitrap LC/MS. A. Linearity check for representative BMPs and PGs, corresponding to Fig. 3B. B. Linearity check for representative BMPs and PGs, corresponding to Fig. 4A. Linearity checks from pooled quality controls (QCs) of samples from corresponding figures. Linearity check R^2^ by simple linear regression.

**Figure S4.**
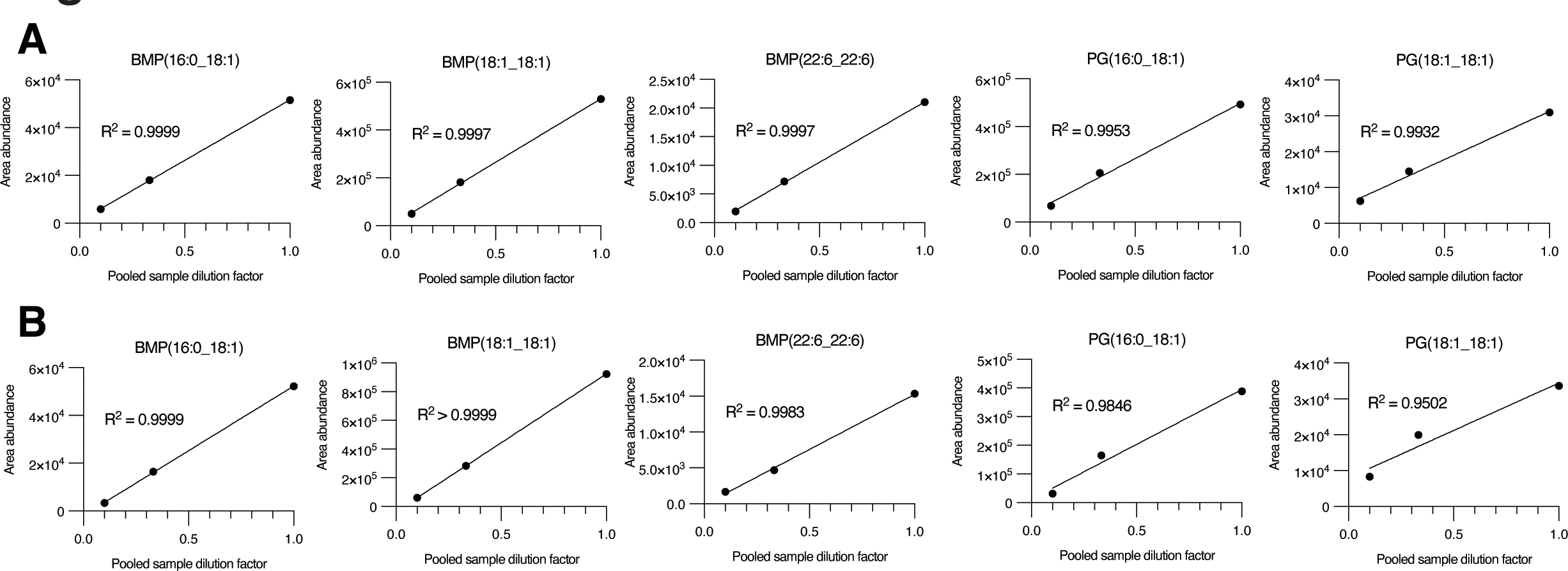
Linearity quality control check for BMP and PG profiling by QQQ LC/MS. A. Linearity check for representative BMPs and PGs, corresponding to Fig. 3C. B. Linearity check for representative BMPs and PGs, corresponding to Fig. 4B. Linearity checks from pooled quality controls (QCs) of samples from corresponding figures. Linearity check R^2^ by simple linear regression.

**Figure S5.**
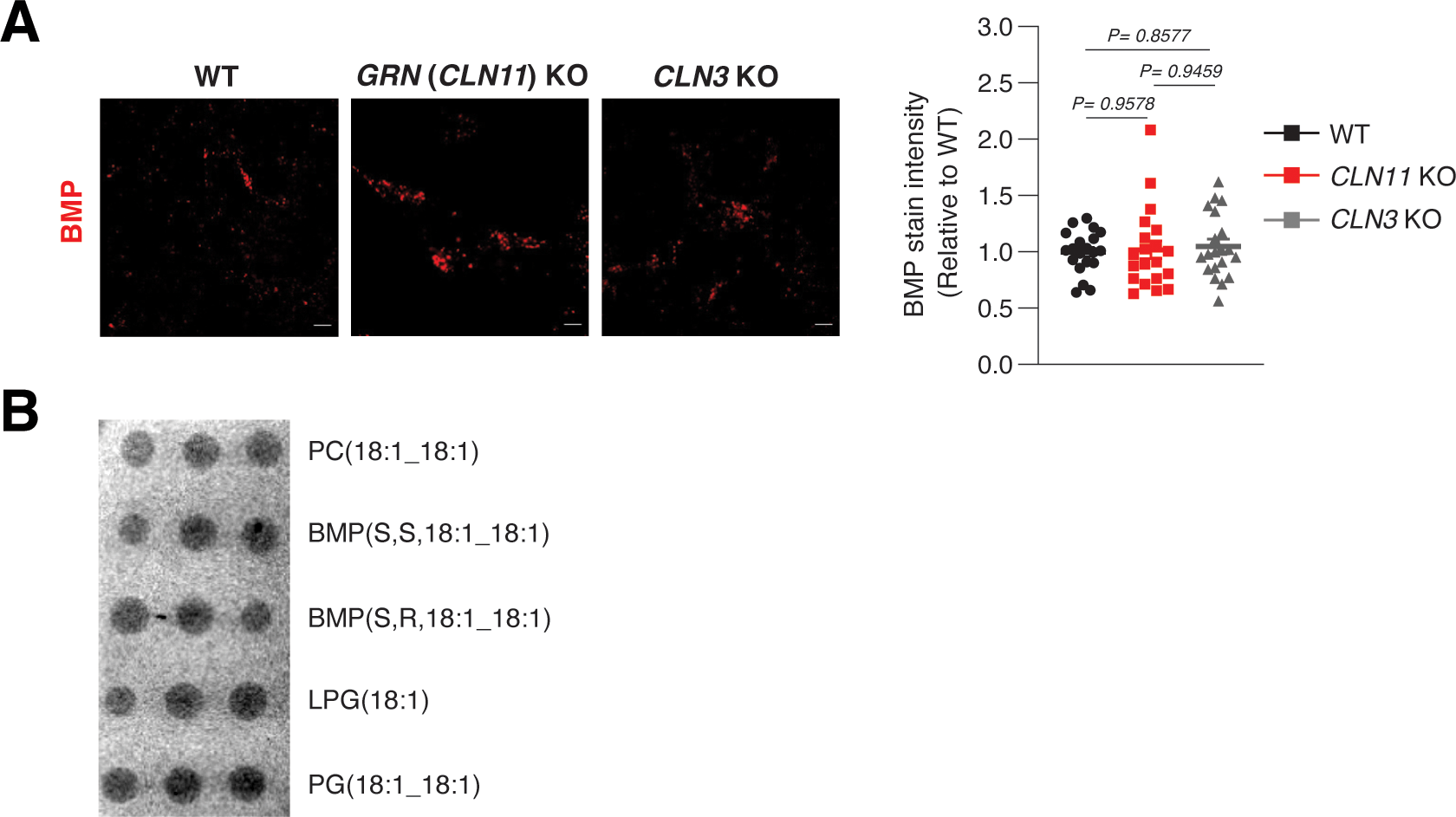
Comparison of BMP profiling by LC/MS and anti-BMP staining. A. BMP levels in *CLN11* KO and *CLN3* KO quantified by anti-BMP antibody. B. Lipid dot blot for indicated phospholipids detected by anti-BMP staining. Anti-BMP staining in WT, *GRN* (*CLN11*) KO, and *CLN3* KO HEK293T cells. Images presented with scale bar = 5 µM. Data is presented as mean ± SEM (n = 20). P-values by one-way ANOVA with post-hoc Tukey’s HSD test. Lipid dot blot is presented with n = 3. PC = phosphatidylcholine; LPG =lysophosphatidylglycerol.

